# Functional characterisation of bicarbonate transporters from the cyanobacterial SbtA2 family and subsequent expression in tobacco

**DOI:** 10.1101/2025.11.03.686429

**Authors:** Loraine M. Rourke, Caitlin S. Byrt, Benedict M. Long, G. Dean Price

## Abstract

Cyanobacteria rely on bicarbonate (HCO_3_^-^) as the primary inorganic carbon (Ci) source for photosynthesis in aquatic environments. To use of this Ci source, cyanobacteria employ CO_2_ concentrating mechanisms (CCMs) that elevate cytoplasmic HCO_3_^-^ via plasma membrane transporters, enhancing carboxylation by carboxysomal Rubisco. The sodium-dependent SbtA1 transporter family is well-characterized in freshwater cyanobacteria, but the related SbtA2 family, prevalent in marine α-cyanobacteria, remains uncharacterised. Here, we report functional characterisation of SbtA2 homologues from marine *Synechococcus* spp., which exhibit high Ci uptake flux with apparent chloride dependence and intermediate HCO_3_^-^ affinity (Km ≈ 150 µM), when assessed in *E. coli*. SbtA2 achieved internal Ci accumulation up to 24 mM within 30 seconds. Co-expression with the putative regulator SbtB2 reduced uptake activity, suggesting a regulatory role for this protein. These findings indicate that SbtA2 transporters contribute significantly to carbon acquisition in marine cyanobacteria. Given potential to enhance CO_2_ supply to Rubisco in C_3_ plants, we targeted SbtA2 to the tobacco chloroplast inner envelope membrane; however, this did not improve photosynthesis or growth. Our results highlight the functional diversity of cyanobacterial Ci transporters and suggest that additional components may be required for effective transfer of such systems into plant chloroplasts.

**Highlight:** SbtA2 bicarbonate transporters from some marine α-cyanobacteria are high flux transporters with an unusual chloride-dependence and show intermediate uptake affinity when expressed in *E. coli*. Initial attempts failed to demonstrate enhanced photosynthesis when directed to tobacco chloroplasts.

## Introduction

Cyanobacteria are single-celled, photosynthetic organisms of global ecological importance due to their significant contribution to CO_2_ fixation into the biosphere (Field *et al*., 1998; Flombaum *et al*., 2013). Their acquisition of carbon is primarily driven by the enzyme Rubisco, which fixes CO_2_ as part of the Calvin -Benson -Bassham (CBB) cycle for use in cellular metabolism (Badger *et al*., 2002; Nguyen *et al*., 2025a). While CO_2_ is readily able to permeate cellular membranes, the most abundant form of inorganic carbon (Ci; CO_2_, CO_3_^2-^, and HCO_3_^-^) available for photosynthesis in aquatic environments is the relatively membrane-impermeable bicarbonate ion (HCO_3_^-^; Falkowski and Raven, 2007). Cyanobacteria and other aquatic photoautotrophs have evolved CCMs to enable the use of this Ci source as a means to elevate CO_2_ around Rubisco and maximise carboxylation (Badger *et al*., 2006; Price *et al*., 2008). Cyanobacterial CCMs are highly efficient bipartite systems, utilizing Ci transporters to enable active accumulation of HCO_3_^-^ in the cytoplasm, and HCO_3_^-^-permeable micro-compartments called carboxysomes in which Rubisco is encapsulated (Rae *et al*., 2013a; Nguyen *et al*., 2025b). Within carboxysomes, a specialized carboxysomal carbonic anhydrase (CA) equilibrates HCO_3_^-^ with CO_2_ (Reinhold *et al*., 1991; Pulsford *et al*., 2024), thus increasing CO_2_ available for carboxylation (Long *et al*., 2021; Rae *et al*., 2013b).

There are two main classifications of cyanobacteria based on the form of Rubisco they contain (Badger and Bek, 2008). Using Form-IB Rubisco, β-cyanobacteria are frequently found in smaller freshwater bodies and estuarine habitats, whilst the more abundant α-cyanobacteria (containing Form-IA Rubisco) are widespread in both oceanic and freshwater environments (Badger *et al*., 2002; Cabello-Yeves *et al*., 2022). Some α-cyanobacteria are referred to as transition strains and show evidence of genetic inheritance of CCM components by horizontal transfer, often derived from β-cyanobacteria (Rae *et al*., 2011).

The α-cyanobacteria are thought to have a significant impact on global CO_2_ acquisition into the biosphere due to their relative abundance (Field *et al*., 1998; Cabello-Yeves *et al*., 2022) and high rate of CO_2_ fixation (Bar-On and Milo, 2019). Estimates suggest as much as 25% of marine primary productivity can be attributed to the predominant α-cyanobacterial genera, marine *Synechococcus* spp. and *Prochlorococcus* spp. (Flombaum *et al*., 2013).

Underpinning CCM function across these phylogenies is a suite of Ci transport systems that support cellular HCO_3_^-^ elevation. Currently, there are five Ci transport systems involved in cyanobacterial CCMs that have been characterised: two CO_2_-to-HCO_3_^-^ conversion complexes bound to thylakoid membranes, and three families of plasma membrane HCO_3_^-^ transporters (Price *et al*., 2008; Long *et al*., 2016). The HCO_3_^-^ transporters include the constitutively expressed, high affinity/low flux ABC-type transporter BCT1 (Omata *et al*., 1999; Rottet *et al*., 2024), the inducible high-affinity/low flux transporter SbtA1 (Shibata *et al*., 2002; Du *et al*., 2014), and the low affinity/medium-high flux transporter BicA (Price *et al*., 2004). Directional conversion of CO_2_ -to -HCO_3_^-^ is performed by specialized Type 1 NDH complexes on the thylakoid membranes that use the electron transport chain and specialised CupA/CupB (Walker *et al*., 2024; Maeda *et al*., 2002). Both HCO_3_^-^ and CO_2_ uptake systems are found in most cyanobacterial species, with the exception of *Prochlorococcus* spp., which generally possess only distant homologues of BicA and SbtA (Rae *et al*., 2011).

Human nutrition relies heavily on C_3_ crop plants (Salesse-Smith *et al*., 2025). However, under current atmospheric conditions, C_3_ plants have inefficient photosynthetic processes compared to most other photosynthetic organisms (for instance C_4_ plants and phytoplankton) due to a lack of a CCM. CCMs act to increase the concentration of CO_2_ at the site of Rubisco, the primary enzyme responsible for carboxylation in photosynthetic organisms (Erb and Zarzycki, 2018). One approach to improve the photosynthetic efficiency of these plants is to transplant a CCM from cyanobacteria into the chloroplast in order to improve CO_2_ delivery to Rubisco (Price *et al*., 2011). However, there is a knowledge gap in the mechanistic and regulatory understanding of cyanobacterial HCO_3_^-^ transporters that needs to be filled before such an ambitious goal can be achieved (Rottet *et al*., 2021). Modelling indicates that incorporating a carboxysome in the chloroplast stroma, and expression of HCO_3_^-^ transporters in the inner envelope membrane (IEM) of the chloroplast, could increase plant productivity by 36-60% (McGrath and Long, 2014). The expression of a HCO_3_^-^ transporter in the chloroplast IEM alone is also likely to improve photosynthetic efficiency in plants by enhancing net CO_2_ supply to Rubisco (Price *et al*., 2011; McGrath and Long, 2014). HCO_3_^-^ transport proteins have been expressed in the chloroplastic IEM, however there has been little evidence of active HCO_3_^-^ uptake in the chloroplast so far (Rottet *et al*., 2024; Rottet *et al*., 2021; Forster *et al*., 2023; Nölke *et al*., 2019; Atkinson *et al*., 2016; Pengelly *et al*., 2014). While it is unknown as to why this is the case, it is possibly due to a lack of activation mechanisms for foreign transport proteins or incorrect conditions to drive transporter function (Rottet *et al*., 2021). For instance, for Na^+^-dependent HCO_3_^-^ transporters such as SbtA1 and BicA, the Na^+^ concentration in the cytoplasm (and therefore the cytoplasm:chloroplast stromal Na^+^ gradient) may not be optimal for function *in planta*. This demonstrates the need to further understand the function of these HCO_3_^-^ transporters and identify different classes of transporters that could be adapted for use in C_3_ chloroplasts.

In this study, we focus on the previously undescribed HCO_3_^-^ transporter gene family *sbtA2* (<u>s</u>odium-dependent <u>b</u>icarbonate transporter A2) which is largely found in the α-cyanobacteria (Figure 1; Cabello-Yeves *et al*., 2022; Rae *et al*., 2011) and has distant protein homology to the previously characterized sodium-dependent *sbtA1* family (Figure S1; Price *et al*., 2008; Rae *et al*., 2011; Du *et al*., 2014).(Price *et al*., 2008; Rae *et al*., 2011; Du *et al*., 2014). Similar to *sbtA1*, the *sbtA2* gene is typically co-located in cyanobacterial genomes with *sbtB2* (Figure S5A-D), coding for a putative regulatory protein SbtB2, homologous to the regulator SbtB1 (Du *et al*., 2014; Förster *et al*., 2023; Kaczmarski *et al*., 2019).

**Figure 1:**
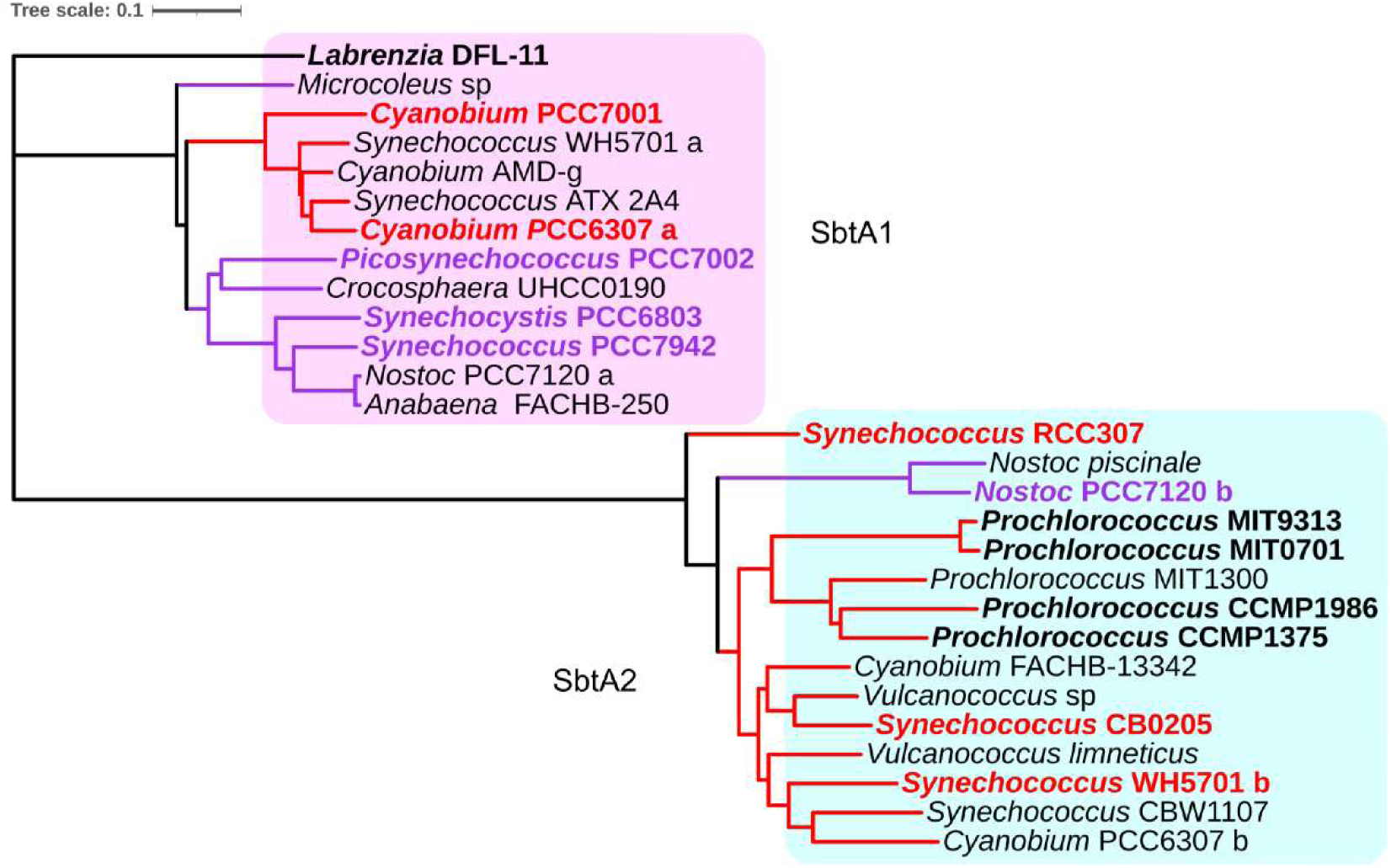
Phylogenetic tree showing the relative protein relationships between members of the SbtA family across cyanobacteria. The primary sequence alignment, and derived tree, were prepared using Geneious Prime software and iTol (Letunic and Bork, 2024). The SbtA2 family is highlighted in blue and the SbtA1 family in pink. The internal size marker indicates 0.1 substitutions per residue. Red branches indicate α-cyanobacteria and purple branches indicate β-cyanobacteria. Coloured bold text indicates that HCO3^-^ uptake activity has been assessed with those and in black bold text demonstrating no function in E. coli (this study and Du *et al*., 2014). The outgroup SbtA1 from *Labrenzia* DFL-11, an α-proteobacterium found in close association with dinoflagellates of the *Alexandrium* genus (Biebl *et al*., 2007), was previously assessed for function by (Du *et al*., 2014).

Here we characterize the function of several members of the SbtA2 family from α-cyanobacteria in a heterologous bacterial expression system. Genes encoding SbtA2 and SbtB2 from marine α-cyanobacterial species were cloned individually, or in tandem, and expressed in a high-CO_2_-requiring (HCR) *E. coli* strain, CAfree (Rottet *et al*., 2024; Forster *et al*., 2023; Desmarais *et al*., 2019). SbtA2 homologues that rescued CAfree at ambient CO_2_ were further characterized by measuring their HCO_3_^-^ uptake rates, counter-ion dependency and affinity for HCO_3_^-^. We also assessed the hypothesis that their associated SbtB2 protein acts as a regulator of SbtA2 function in this system. Our results show that like SbtA1:B1, SbtB2 appears to be capable of limiting the activity of its SbtA2 counterpart in *E. coli*. We provide focused analysis of SbtA2 function from *Synechococcus* WH5701 (hereafter SbtA2-WH5701), revealing a potentially different function from its SbtA1 homologue. We then expressed SbtA2 in the chloroplast IEM of *Nicotiana tabacum* to assess its capacity to improve the photosynthetic efficiency of the plants by increasing CO_2_ supply to Rubisco.

## Materials and Methods

### Molecular cloning of SbtA2 genes

Genes encoding *sbtAB2* from *Synechococcus* WH5701 (accession WP_006170362.1, WP_006170363.1), *Synechococcus* RCC307 (NBQ36408.1, CAK27720.1) and *Synechococcus* CB0205 (WP_010316543.1, WP_010316542.1), *Prochlorococcus* MIT0701 (WP_036914441.1, WP_036914438.1) and *Prochlorococcus marinus* MIT9313 (WP_011130583.1, WP_011130582.1), were synthesized by Genscript (USA) and cloned individually or in tandem under the control of a lac inducible promoter in vectors pCk1 (Kanamycin resistance) or pCsA (Spectinomycin resistance), using type-IIS restriction enzyme cloning that use key features of Golden Gate cloning and Loop Assembly (Pollak *et al*., 2019; Engler *et al*., 2014). In the absence of a suitable antibody for SbtA2, despite numerous attempts to make one, internal epitope tags were introduced to SbtA2 in the cytoplasmic loop between membrane helices five and six of SbtA2 (Table S3) using the same cloning methods.

### CAfree growth complementation assay

A double CA knockout strain of *E. coli* (CAfree; *Δcan ΔcynT*) described by Desmarais et al. (2019), which is unable to grow at air levels of CO_2_, was used to screen for HCO_3_^-^ transport. Cultures of CAfree expressing members of SbtA2 were grown in Luria-Bertani (LB) broth at 37°C in air supplemented with 4% CO_2_ for 18 hours, then diluted 1:5 and grown for an additional hour. The density of the cultures was standardized based on their absorbance (OD_600_ = 0.2) and serially diluted before spotting 10 µL onto LB containing 1.5% agar and appropriate antibiotics (kanamycin – 50 µg/mL or spectinomycin – 100 µg/mL) and 100 µM isopropyl β-D-1-thiogalactopyranoside (IPTG). Plates were incubated overnight at 37°C in air levels (0.04%) and elevated (4%) CO_2_ to test for complementation.

### CAfree growth assay

Cultures of CAfree expressing each form of SbtA2 were incubated in LB supplemented with antibiotics (as for the complementation assay) with shaking for 18 h at 37 °C in CO_2_-supplemented air (4% CO_2_). These were diluted 1:5 and grown for approximately 60-90 minutes to achieve log phase growth. Cultures were diluted 1:10 into final culture volumes of 200 µL and were grown at 37 °C in 96 well plates in a plate reader (BMG Labtech) under air and CO_2_-supplemented-air (4% CO_2_) conditions in LB supplemented with antibiotics and IPTG. Growth was measured by taking automated absorbance (OD_600_) readings at 10-minute intervals for 20 h.

### Inorganic carbon uptake assay

Inorganic carbon uptake was examined in *E. coli* (DH5α strain) expressing *sbtAB2* gene constructs in pCk1 or pCsA (and their derivatives) along with appropriate controls. Starter cultures were grown overnight from a standard glycerol stock in LB, this culture was then diluted 1:5 in LB and grown at 37°C for approximately one hour until absorbance readings (OD_600_) reached 0.6. Expression was induced by the addition of IPTG to a final concentration of 1 mM, and cultures were grown for an additional 2.5 h at 37°C with shaking. Culture density was measured, and cell volumes adjusted to a standard cell density for each experiment with an approximate OD_600_ 1.0-1.2. Cells (∼2 mL) were depleted of Ci by washing twice with 1 mL of CO_2_-free buffer (20 mM BisTrisPropane-H_2_SO_4_, pH 7.5, salts as required; sparged with N_2_ gas) after centrifugation (20,238 × *g* for 30 s). NaH^14^CO_3_ (∼60 MBq/mmol) was added to the cells to a final concentration of 0.5 mM and 200 µL of this suspension was overlayed onto 50 µL silicone oil mix AR200:20 3.5:4, above a 20 µL kill solution (3 M sodium hydroxide, 50% methanol) in a 400 µL polypropylene tube (Beckman Coulter). After 30 second incubation the NaH^14^CO_3_-cell mix was immediately centrifuged for 20 seconds to separate the cells with incorporated ^14^C from free NaH^14^CO_3_ which remains in the supernatant. Tubes were immediately placed in an aluminium block on dry ice, before clipping off the base of the tube containing the cell pellet into scintillation vials containing 400 µL 2 M NaOH. Vials were vortexed well and 2 mL UltimaGold Scintillation cocktail (Perkin Elmer) added before being counted using a Hidex Scintillation counter for 2 minutes. Variations of assay buffer pH, including the concentration of Ci, acid used to pH the buffer, ions added to the uptake buffer and how the NaH^14^CO_3_ stock was diluted were also used.

To characterise the kinetics of SbtA2 transporters, Ci uptake rates were measured over a range of Ci concentrations to cells expressing each SbtA2. The resulting data were analysed using Michalis-Menten non-linear regression with the statistical fit to assess if the transport kinetics of SbtA2 meets the criteria for active transport.

### Determination of SbtA2 cellular localization in *E. coli*

Overnight *E. coli* (DH5α) cultures expressing SbtA2 forms carrying an internal His×6 tag were grown and induced as per Ci uptake assay. Cells were harvested at 4,800 × *g* for 5 minutes, washed in lysis buffer (50 mM HEPES-KOH pH 8.0, 100 mM NaCl, 5% glycerol), and left to incubate with ∼30 kU of rLysozyme (Merck Millipore) for 30 minutes at 37°C with gentle shaking. Cells were broken using approximately 200 µL silica beads using a mini beadbeater-16 homogeniser (BioSpec Products) for 5 minutes. Cell debris and beads were removed by centrifugation (20,238 × *g*, 1 minute at room temperature). Supernatants were transferred to clean tubes and membrane enriched fractions collected by centrifugation at 16,700 × *g* in an Eppendorf bench top centrifuge for 30 minutes at 4 °C. Pellets were resuspended in a buffer containing 50 mM HEPES pH 8.0, 100 mM NaCl, 25 mM imidazole, and 1% *n-*Dodecyl*-β-D-* maltoside (DDM) for 30 minutes on ice to solubilise membrane proteins. Insoluble debris was removed by centrifugation, 16,700 × *g* at 4 °C for 20 minutes, and the supernatant taken was used as the membrane enriched fraction.

### SDS-PAGE and western blot

Proteins were separated by SDS PAGE (NuPAGE, 4-12% Bis-Tris mini protein gel, Thermo Fisher Scientific) and subsequently transferred on to PVDF membrane for 45 minutes (BioRad Transblot Turbo). The membrane was blocked with 5% skim milk powder and probed for His×6 or HA tagged proteins using mouse-anti-His×6 or mouse-anti-HA antibodies (and goat-anti-mouse AP-conjugated secondary antibody; Table S5), and developed with Attophos (Promega, USA) prior to imaging on a BioRad Chemidoc MP system as previously detailed (Long *et al*., 2018).

## Plant methods

### Nuclear transformation

#### *Agrobacterium* transformation

*Agrobacterium tumefaciens* (strain GV3101) was transformed with 1µg of pDNA by electroporation, and recovered with 0.5 mL LB at 28°C for 2 h before plating on LB-agar supplemented with 30µg/mL kanamycin and 50 µg/mL rifampicin and incubated for ∼48 h at 28°C.

#### *N. tabacum* transformation

*Agrobacterium* harbouring plasmid of interest was cultured in LB supplemented with 30 µg/mL kanamycin and 50 µg/mL rifampicin at 28°C for 48 h. Cells (∼20 mL) were harvested by centrifugation for 10 minutes at 1650 × *g*. Cells were suspended in resuspension medium, and left at room temperature for 1 h. Sterile tobacco leaves were cut from plantlets growing in tissue culture into 0.4 cm^2^ pieces and placed adaxial side down on co-cultivation plates with 5 mL of *Agrobacterium* in resuspension medium and left for 5 minutes, before transferring plant pieces to fresh co-cultivation plates. Plates were sealed and left in a controlled temperature growth room at 25°C with 16:8 h photoperiod.

After 48 h leaf pieces were transferred to regeneration plates. When shoots appeared, callus with shoots were transferred to fresh regeneration plates and returned to the growth room. When the shoots became big enough to excise they were removed from the leaf pieces and transferred to rooting plates. Once roots had formed, plantlets were transferred to soil (6L of commercial seed raising mix with 2-3 cups of vermiculite and ∼25-30 mL of Osmocote) in 8×8 cm^2^ punnets in growth chamber.

#### Growth chamber conditions

Temperature was controlled at 25°C daytime/ 20°C night time, relative humidity 60%, light intensity of 400 µmol s^-1^ m^-2^, daylength 0700-2300 h. After 1-2 weeks, leaf samples (1 cm^2^ discs) were taken for determination of the hygromycin resistance gene insert number using droplet digital PCR (iDNA Genetics Ltd, Norwich, UK). Only plants with one insert were continued to the next stages. Leaves were also sampled at this time for immunoblot analysis to assess the expression levels of the transgene. When necessary, plants were transferred to 4.5 L pots filled with steam-sterilised Martins Mix supplemented with 7 g/L Osmocote and moved to the glasshouse to reach maturity prior to seed harvest.

For plant growth in glasshouses, pots were placed on benches ∼ 1 m off the ground and temperature was controlled to 25°C daytime/20°C night by commercial air-conditioning.

### Plant height, fresh and dry weight determination

The height of plants was measured each day from the base of the stem to the base of the apical bud. At 63 days post cotyledon emergence (pce), the above-ground tissue of plants was harvested by cutting at the base of the stem and the entire plant was placed into a paper bag and weighed. The weight of the paper bag was subtracted to determine fresh weight. Plants were then placed in 80°C oven for seven days to dry the plant tissue. The weight of the tissue and bag was recorded, and the weight of the bag subtracted to obtain the dry biomass weight.

### Δ^13^C determination of plant biomass

Leaf samples (0.5 cm diam discs) were harvested following gas exchange measurements (30 ± 5 cm in height). Samples were placed in 1.5 mL tubes and dried at 80°C for 24 h. The tubes were capped and Δ^13^C determination was performed by mass spectrometer at RSB, ANU (Förster *et al*., 2023)

### Leaf gas exchange measurements

Using LI-6800 (LICOR) with a 6 cm^2^ aperture, fully expanded leaves of plants with similar size (30 cm ± 5cm) and leaf number were equilibrated to the room for ∼20 minutes before gas exchange measurements. Leaves were equilibrated at 400 µmol mol⁻¹ CO_2_ for the reference for 10-20 minutes before starting CO_2_ response curves over a series of CO_2_ concentrations at two-minute intervals (400, 200, 100, 50,150, 250, 400, 600, 800, 1200, 400 µmol mol⁻¹ for reference) at a leaf temperature of 25°C, 55% relative humidity, and illumination of 1500 µmol s^-1^ m^-2^.

A/C_int_ curves were generated from the data by plotting Assimilation rate (A) against intercellular CO_2_ concentration (C_int_) as calculated in the LI-6800 output.

#### Estimation of CO_2_ compensation point

The CO_2_ compensation point (the intercellular CO_2_ concentration where net CO_2_ assimilation rate is 0 µmol m^-2^ s^-1^) was determined from the A/C_int_ curves. The linear part of the curve (0-300 µmol mol^-1^ intercellular C_int_) was used to determine the compensation point.

### Chloroplast membrane enrichment

Chloroplast membrane-enriched protein fractions were isolated from two leaf discs (12 mm^2^) that were snap frozen when sampled and stored at -80°C. Samples were ground using Dounce homogenisers with 1 mL of lysis buffer (50 mM EPPS pH 8, 5 mM MgCl_2_, 2 mM DTT, 1 mM Na-diethyl-dithiocarbamate, 0.01g/mL polyvinylpyrrolidone [PVPP]) and 10 µL plant protease inhibitor cocktail (Cat # P9599, Merck Life Science, USA). The suspension was transferred to a 1.5 mL tube, on ice and centrifuged for 30 s at 20,238 × *g* to remove large debris and PVPP. The supernatant was transferred to a new 1.5mL tube and centrifuged for 15 minutes at 20,238 × *g* at 4°C. The aqueous supernatant was discarded before resuspending the pellet in wash buffer (50 mM EPPS pH 8, 5 mM MgCl_2_, 2 mM DTT, 1 mM Na-diethyl-dithiocarbamate, 200 mM NaCl). Centrifugation and washing were repeated twice. All traces of supernatant were removed, and the pellet was resuspended in 50 µL of 2 × resuspension buffer (4% SDS, 125 mM tris-HCl pH 8.0, 1 mM EDTA pH 8.0) and left at 4°C overnight. The membrane-enriched suspension was centrifuged for 1 minute at 20,238 × *g* at 4°C and the soluble fraction was transferred to a new tube.

Total protein in the samples was determined using the detergent compatible Bio-Rad DC Protein Assay kit, by diluting the samples 1:10 in 2 × resuspension buffer. Sample (5 µL) or BSA standard was added in triplicate in a 96-well plate, and the kit used as per instruction protocol. Samples were standardised based on total protein values of the samples before being loaded onto SDS PAGE for immunoblot analysis.

## Results

Based on phylogenetic analysis of multiple protein sequence alignments of SbtA transporter proteins, the well characterised SbtA1 HCO_3_^-^ transporters (Du *et al*., 2014), largely present in freshwater and estuarine β-cyanobacteria, are distinct from homologues found in α-cyanobacteria (Figure 1; Cabello-Yeves *et al*., 2022; Rae *et al*., 2011; Du *et al*., 2014). On this basis, we refer here to the former group as SbtA1, and the latter as SbtA2 (Rae *et al*., 2011). As an example of the phylogenetic distance between SbtA1 and SbtA2 members, sequence identity of SbtA1 and SbtA2 from *Synechococcus* WH5701 is approximately 25% (Figure S1A).

### Heterologous expression and bicarbonate transport function of SbtA2 homologues

To evaluate the HCO_3_^-^ transport function of SbtA2 homologues, genes from members of the SbtA2 family (from marine α-cyanobacterial species *Synechococcus* WH5701, RCC307, CB0205, and *Prochlorococcus* MIT0701, and MIT9313) were expressed in the *E. coli* strain CAfree (Desmarais *et al*., 2019) under the control of the IPTG-inducible *lacI^q^-trc* hybrid promoter (de Lorenzo *et al*., 1993).

The CAfree strain of *E. coli* lacks both CA genes *cynT* and *can* (Desmarais *et al*., 2019) and is therefore incapable of enzymatic HCO_3_^-^ to-CO_2_ interconversion within the cytoplasm. This renders CAfree reliant on exogenous CO_2_ entry and slow spontaneous chemical conversion to HCO_3_^-^ as a carboxylation substrate to supply anaplerotic pathways in the cell (Merlin *et al*., 2003). CAfree is unable to grow under atmospheric CO_2_ concentrations (∼0.04% CO_2_) as the rate of HCO_3_^-^ supply is too slow to support these pathways. The subsequent phenotype is a dependency on either high-CO_2_ for growth, due to high relative rates of spontaneous HCO_3_^-^ production within the cell, or heterologous expression of either a functional CA (Pulsford *et al*., 2024) or HCO_3_^-^ transporter (Rottet *et al*., 2024; Forster *et al*., 2023) to grow at air levels of CO_2_. In our hands we have seen no reversion of this cell line under standard experimental conditions.

Of the five SbtA2 family members assessed (chosen as genes with moderate GC content that therefore did not require codon optimisation), three supported growth of CAfree at ambient CO_2_ (Figure 2A, S2A). These were from *Synechococcus* spp. WH5701, RCC307 and CB0205; referred to hereafter as SbtA2-WH5701, SbtA2-RCC307 and SbtA2-CB0205 respectively. Neither of the SbtA2 homologues from *Prochlorococcus* spp. (MIT0701 and MIT9313) showed function in CAfree at ambient CO_2_ under the complementation conditions assessed (Figure 2A). Functional SbtA2 proteins were expressed in *E. coli* strain DH5α and characterised by radiolabelled NaH^14^CO_3_ uptake assays using the silicone oil centrifugation-filtration method to determine their HCO_3_^-^ transport activity (Rottet *et al*., 2024; Du *et al*., 2014; Forster *et al*., 2023).

**Figure 2:**
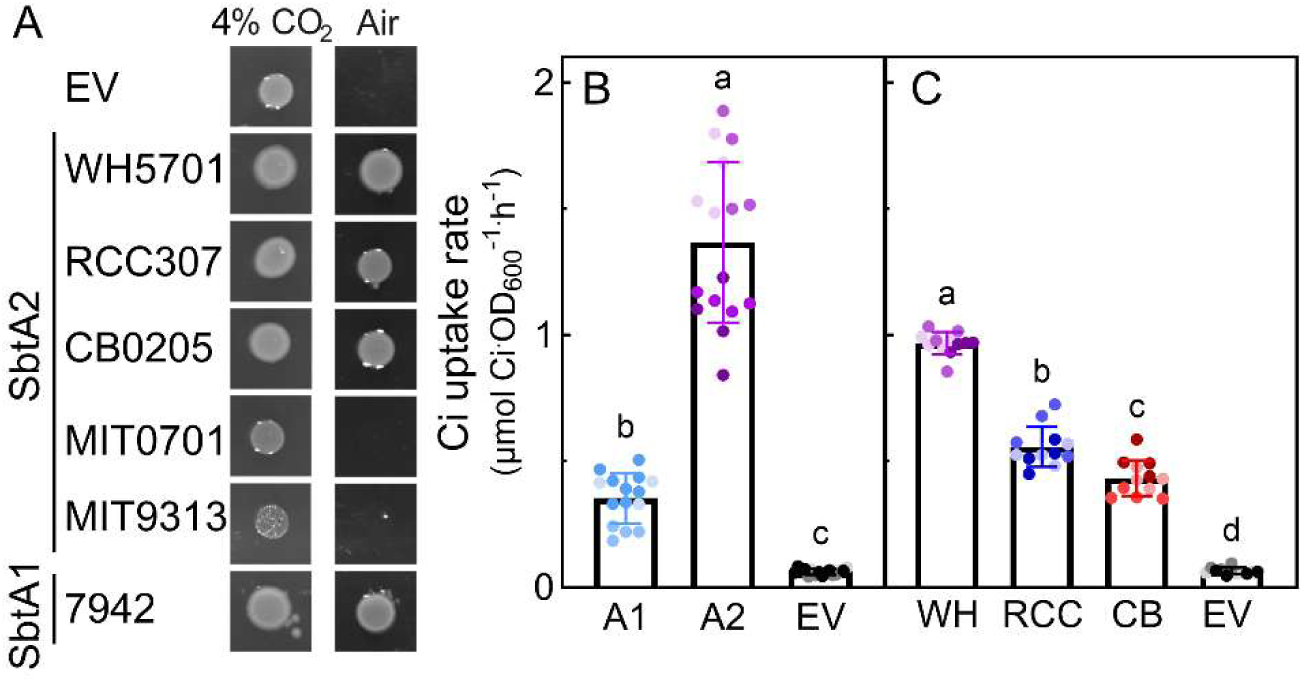
The HCO ^-^ transport function of SbtA2 was assessed using complementation assays and HCO ^-^ uptake assays. **(A)** Cultures of CAfree E. coli expressing SbtA2 homologues were adjusted to the same density prior to being spotted onto LB-agar containing 100 µM IPTG and grown at ambient (0.04%) or 4% CO2 for ∼20 h. The growth of SbtA2 members (Synechococcus WH5701, RCC307, CB0205 and Prochlorococcus MIT0701 and MIT9313) were compared to a negative control (empty vector; EV) and a positive control (SbtA1 from Synechococcus elongatus PCC7942; SbtA1-7942). **(B)** The HCO3^-^ uptake activity of SbtA2-WH5701 (A2), SbtA1 from Synechococcus PCC7942 (SbtA1-7942; A1) and empty vector (EV). n=12 from 3 biological replicates. **(C)** HCO3^-^ uptake activity of SbtA2 from Synechococcus WH5701 (WH), RCC307 (RCC) and CB0205 (CB). n=12 from 3 biological replicates. HCO3^-^ uptake assay buffer composition: BTP - H2SO4 pH 7.5 + 20 mM NaCl. Different letters indicate significantly different means when analysed by one-way ANOVA (P<0.05).

Previous analysis of members of the SbtA1 family revealed they were Na^+-^-dependent (Du *et al*., 2014). Due to their similarity to the SbtA1 family, SbtA2 function was initially assessed under similar assay conditions used for SbtA1, namely in the presence of 20 mM NaCl (Du *et al*., 2014). Under these conditions, we observed high rates of HCO_3_^-^ transport by SbtA2 (Figure 2B, C), particularly for SbtA2-WH5701 which had significantly higher flux than SbtA1 from *Synechococcus* PCC7942 (SbtA1-7942) when compared on a cell density basis in the presence of 20 mM NaCl (Figure 2B). Typical HCO_3_^-^ uptake rates for SbtA1-7942 were less than 0.5 µmol Ci.OD_600_^-1.^h^-1^ (Figure 2B), whereas SbtA2-WH5701 achieved 1-1.5 µmol Ci.OD_600_^-1.^h^-1^ (Figure 2B, C).

Since SbtA2 was active in the presence of 20 mM NaCl, we hypothesised that SbtA2 might also be Na^+^-dependent. Notably, when SbtA1-7942 was assessed for HCO_3_^-^ uptake in response to Na^+^, this Na^+^-dependent transporter showed very little uptake in buffers with no added NaCl (Figure 3A). However, SbtA2-WH5701 demonstrated elevated rates of HCO_3_^-^ uptake across the range of 0 – 20 mM NaCl (Figure 3A, B), and in the presence of 200 mM NaCl the HCO_3_^-^ uptake rate of SbtA2-WH5701 was as high as 2.5 - 3 µmol Ci.OD_600_^-1.^h^-1^ (Figure 3C).

**Figure 3:**
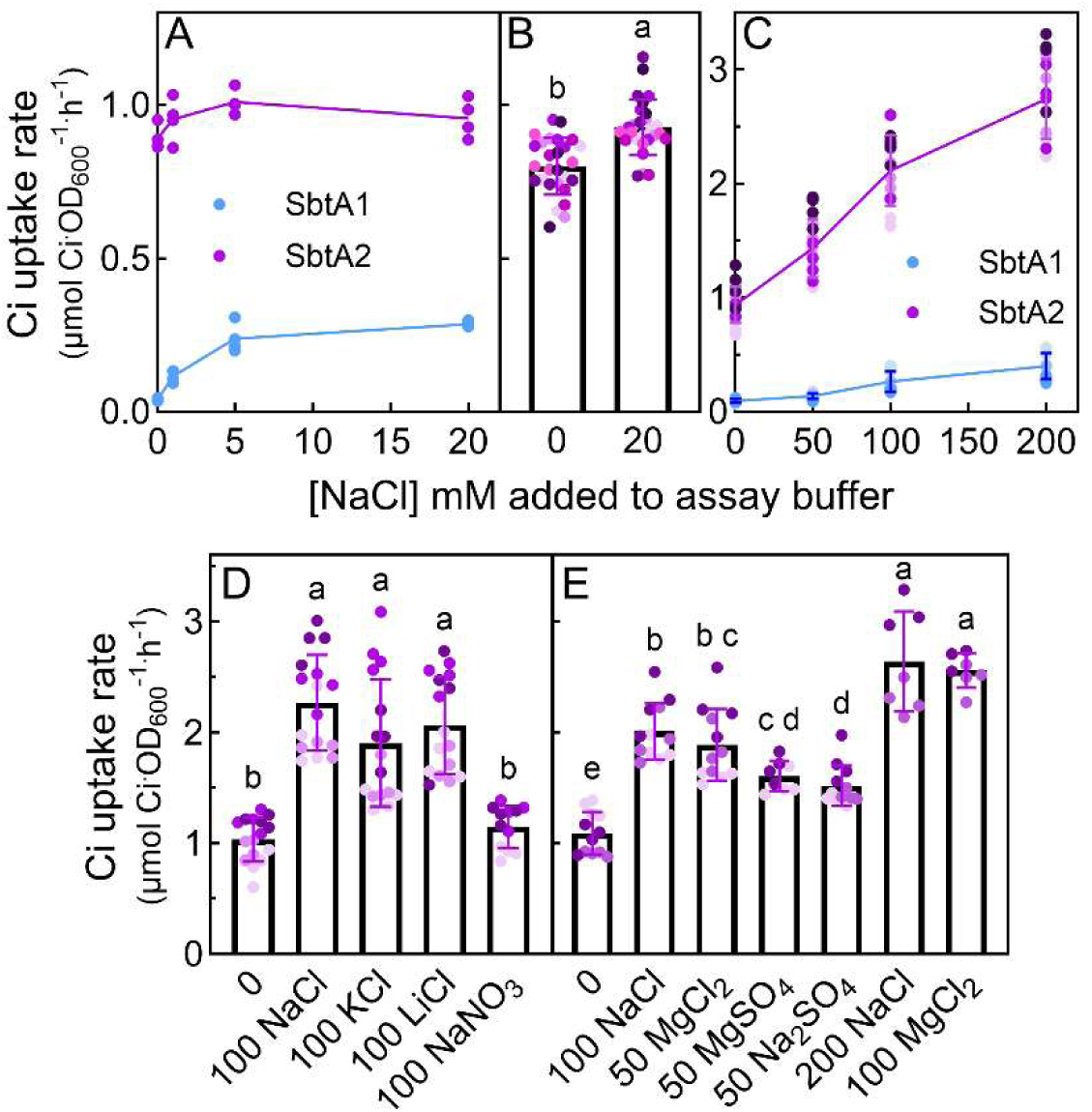
Assessment of the counter-ion driving the HCO3^-^ uptake function of SbtA2. Analysis of ion dependency was determined using HCO3^-^ uptake assays in DH5α strain of E. coli. Values presented have empty vector rates for each condition subtracted. Different letters indicate significantly different means when analysed by one-way ANOVA (P<0.05). **(A)** The HCO3^-^ uptake activity of SbtA2-WH5701 (SbtA2) and SbtA1-7942 (SbtA1) was assessed in the presence of 0, 1, 5 or 20 mM NaCl. Assay buffer composition: 20 mM BTP-H2SO4 pH 7.5 + (0, 1, 5, or 20 mM NaCl). Na^+^ provided by NaH^14^CO3/KHCO3 was ∼20 µM. The connecting line represents the mean; n= 3-4 from 1 biological replicate. **(B)** The HCO3^-^ uptake rates of SbtA2-WH5701 in the presence of 0 and 20 mM NaCl in the assay buffer were compared. Assay buffer composition: 20 mM BTP-H2SO4 pH 7.5 + (0 or 20 mM NaCl). Data presented as mean ± SD; n=27 from 8 biological replicates. **(C)** The HCO3^-^ uptake activity of SbtA2-WH5701 was measured in response to an increase in NaCl concentration in the assay buffer. Uptake assay buffer composition: 20 mM BTP-H2SO4 pH 7.5 + 10 mM K2HPO4 + (0, 50, 100 or 200 mM NaCl). n=11-16 from 3-4 biological replicates. **(D)** The HCO3^-^ uptake activity of SbtA2-WH5701 was assessed in the presence of various salts; no additional NaCl, 100 mM NaCl, 100 mM KCl, 100 mM LiCl and 100 mM NaNO3 added to the assay buffer of 20 mM BTP-H2SO4 pH 7.5. Salt concentrations added to the uptake buffer are in mM. 20 µM Na^+^ was supplied by NaH^14^CO3. Data presented as mean ± SD; n=16-20 from 4-5 biological replicates. **(E)** The HCO3^-^ uptake activity of SbtA2-WH5701 was assessed in response to different salts; no additional NaCl, 100 mM NaCl, 50 mM MgCl2, 50 mM MgSO4 and 50 mM NaSO4, 200 mM NaCl or 100 mM MgCl2 added to the assay buffer of 20 mM BTP-H2SO4 pH 7.5. 20 µM Na^+^ was supplied by NaH^14^CO3. Data presented as mean ± SD; n=8-12 from 2-3 biological replicates.

Based on this result, we then assessed which ion, Na^+^ or Cl^-^, was contributing to the HCO_3_^-^ uptake function by SbtA2, by performing assays in the presence or absence of Na^+^ or Cl^-^ in the uptake assay buffers. SbtA2 HCO_3_^-^ uptake rates were elevated in the presence of NaCl, KCl and LiCl above a control with no added salt, but not in the presence of NaNO_3_ (Figure 3D), revealing that SbtA2 HCO_3_^-^ uptake activity increased in the presence of Cl^-^ and not Na^+^. This observation was further assessed by measuring the HCO_3_^-^ uptake activity of SbtA2-WH5701 in the presence of NaCl, MgCl_2_, Na_2_SO_4_ or MgSO_4_, revealing the possibility of both Cl^-^ or SO_4_^2-^-dependency (Figure 3E).

### SbtA2 effect on *E. coli* Ci pool sizes

Using cell volumes measurements for *E. coli* previously estimated by our group (Du *et al*., 2014), we calculated the internal Ci pool accumulation for a 30 second period from the addition of 0.5 mM Ci to be 5.5 - 8 mM for SbtA2-WH5701, 2 – 3 mM for SbtA1-7942 and 0.2 mM for the empty vector in the presence of 20 mM NaCl. However, in the presence of 200 mM NaCl, the pool size increased to 15 - 23.5 mM for cells carrying SbtA2-WH5701 and 0.3 mM for cells carrying the empty vector (Table 2).

### SbtA2 affinity for HCO_3_^-^

HCO_3_^-^ affinity estimates (K_m_) were calculated to be approximately 150 - 200 µM for all three SbtA2 homologues from *Synechococcus* in the presence of 20 mM NaCl (Figure 4, S4, Table 1 and Table S4). Maximum HCO_3_^-^ uptake rates for SbtA2-WH5701 in the presence of 100 mM NaCl were approximately double that observed at 20 mM NaCl, but the K_m_ for HCO_3_^-^ was unaffected by the increased NaCl concentration (Figure 4 and Table 1).

**Figure 4:**
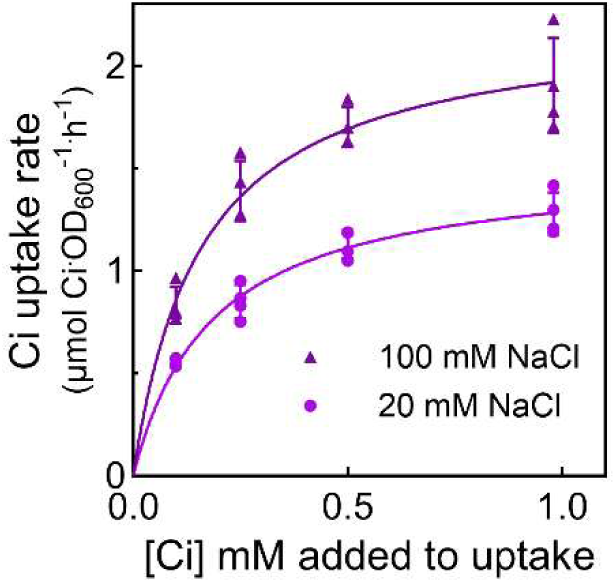
The affinity (Km) for HCO ^-^ was determined for SbtA2 from *Synechococcus* WH5701. The affinity for HCO3^-^ was estimated using Ci uptake assays performed in response to increasing exogenous HCO3^-^ concentrations in E. coli DH5α. The HCO3^-^ affinity of SbtA2-WH5701 was assessed in the presence of 20 mM NaCl or 100 mM NaCl. Assay buffer composition: BTP - H2SO4 pH 7.5 + (20 mM NaCl or 100 mM NaCl). HCO3^-^ uptake rates at each HCO3^-^ concentration had empty vector rates specific for that condition subtracted. Similar trends were observed across experiments although the activity values varied, as a result representative data from single experiments are shown. Individual data values are shown along with their fitted Michaelis-Menten curve which is presented as a solid line. Curve fitting protocols are stated in the materials and methods section.

**Table 1:** The affinity for HCO ^-^ of SbtA2 homologues.

| | WH5701 | WH5701 $\text{Cl}^-$ <sup>a</sup> | RCC307 | CB0205 |
| --- | --- | --- | --- | --- |
| $V_{\max}$ | $1.34 \pm 0.07$ | 2.19 | $0.38 \pm 0.06$ | $0.34 \pm 0.02$ |
| $K_m$ ($\mu\text{M}$) | $146 \pm 33$ | 140 | $147 \pm 16$ | $197 \pm 21$ |

The $K_m$ ( $\mu\text{M}$ ) and $V_{\max}$ ( $\mu\text{mol Ci-OD}_{600}^{-1}\text{h}^{-1}$ ) of the transporters in the presence of 20 mM NaCl in the assay buffer, mean from 3 assays. <sup>a</sup> Results for WH5701 $\text{Cl}^-$ are from a single $\text{HCO}_3^-$ uptake experiment assessing the $K_m$ in of SbtA2-WH5701 in the presence of 100 mM NaCl in the assay buffer.

### The effect of SbtB2 on SbtA2 HCO_3_^-^ uptake activity

Like the *sbtA1* family, *sbtA2* is commonly co-located with *sbtB2* in the genome (Du *et al*., 2014) (Figure S5). SbtB1 acts to down regulate SbtA1 function when expressed in *E. coli* and cyanobacteria preventing SbtA1 activity (Du *et al*., 2014; Förster *et al*., 2023; Fang *et al*., 2021). We therefore assessed whether the SbtB2 protein would affect SbtA2 function in CAfree. Dicistronic constructs of *Synechococcus* SbtA2:B2 with a C-terminal HA-tag on SbtB2 (referred to as SbtA2:B2-HA) were assembled similar to SbtA2 alone, under the control of the *lacI^q^-trc* promoter.

Complementation of CAfree with SbtA2:B2-HA exhibited similar growth to CAfree expressing SbtA2 alone at ambient CO_2_ (Figure 5A, S2B). This suggested that SbtB2-HA does not suppress SbtA2 function in this system. However, NaH^14^CO_3_ uptake assays using the DH5α strain of *E. coli* co-expressing SbtA2-WH5701 and SbtB2-HA-WH5701 (SbtA2:B2-HA-WH5701) exhibited a ∼30% reduction in HCO_3_^-^ uptake activity in comparison to SbtA2 alone (Figure 5B). This suggests SbtB2 has a regulatory role for SbtA2, but not sufficient to completely prevent SbtA2 activity in *E. coli*. To determine if the HA tag on SbtB2 was interfering with the likely capacity for SbtB2 to completely inactivate SbtA2, the HCO_3_^-^ uptake activity of SbtA2:B2-WH5701 devoid of epitope-tags was also assessed. SbtA2:B2-WH5701 showed similar HCO_3_^-^ uptake activity to SbtA2:B2-HA-WH5701 (Figure 5B). Assessment of alternative expression constructs with protein expression under alternative promoters indicated that the relative abundance of SbtA2 and SbtB2 protein influences the activity of SbtA2 (Figure 5C and S5G). Higher expression levels of SbtB2 led to a suppression of SbtA2 activity by up to 90% (Figure 5C and S5G).

**Figure 5:**
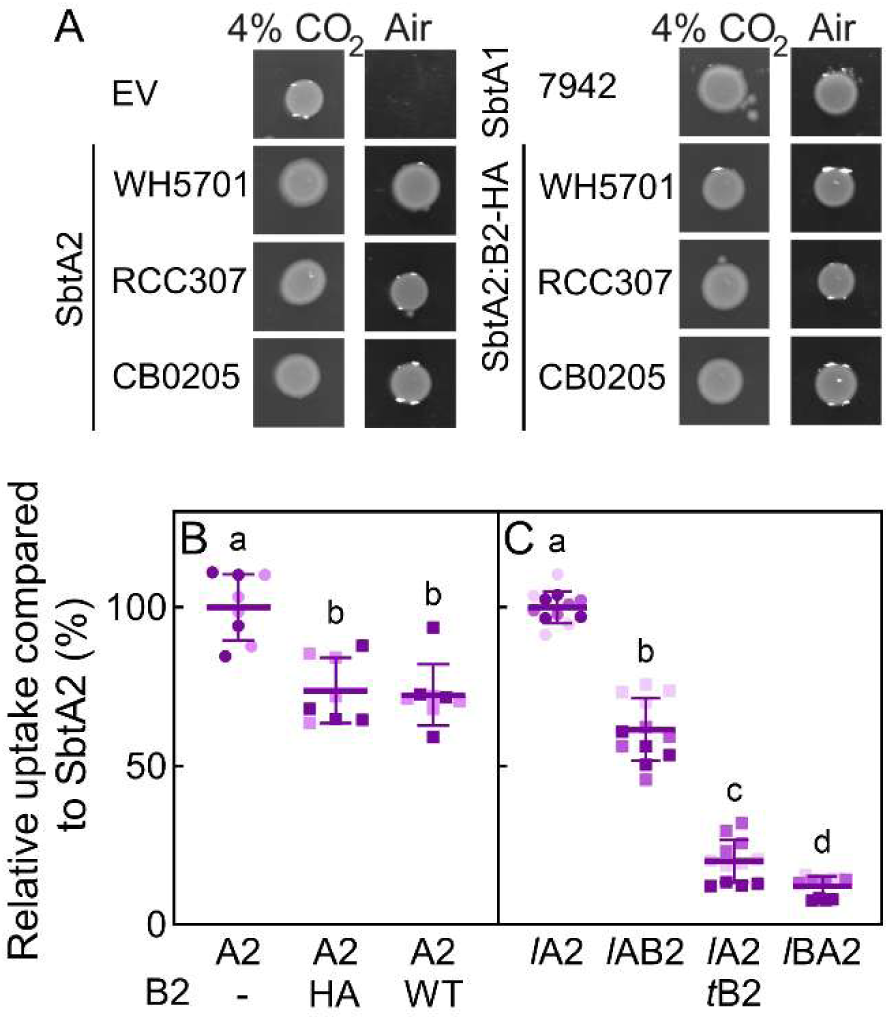
Assessment of the effect of SbtB2 on SbtA2 HCO ^-^ transport function. **(A)** Cultures of CAfree were adjusted to the same density prior to spotting onto LB-agar containing antibiotics and 100 µM IPTG and grown at ambient (0.04%) or 4% CO2 for ∼20 h. The cultures assessed included positive control SbtA1 from *Synechococcus* PCC7942, (SbtA1-7942) SbtA2:B2-HA from *Synechococcus* WH5701, RCC307, and CB0205, a negative control (empty vector; EV) and positive controls (SbtA2-WH5701, SbtA2-RCC307, SbtA2-CB0205). **(B)** The relative HCO3^-^ uptake activity of DH5α *E. coli* carrying SbtA2:B2-HA-WH5701 and SbtA2:B2-WH5701 compared to SbtA2-WH5701 (A2). Assay buffer composition: 20 mM BTP-H2SO4 pH 7.5. Data presented as the mean with empty vector control subtracted ± SD; n=8 from 2 biological replicates. **(C)** The relative HCO3^-^ uptake activity of DH5α *E. coli* carrying *lacI^q^*-SbtA2-H6:B2-HA-WH5701 (*l*AB2), *lacI^q^*-SbtA2-H6:*tet*-B2-HA-WH5701 (*l*A2*t*B2) and *lacI^q^*-SbtB2-HA:A2-H6-WH5701 (*l*BA2) compared to *lacI^q^*-SbtA2-H6-WH5701 (*l*A2). Assay buffer composition: 20 mM BTP-H2SO4 pH 7.5 + 200 mM NaCl. Data presented as the mean ± SD with empty vector controls subtracted; n= 12 from 3 biological replicates.

### Functional characterisation of SbtA2 in *N. tabacum* directed to the chloroplast IEM

In order to assess the potential for SbtA2 to improve net CO_2_ delivery to Rubisco *in planta*, we expressed SbtA2 in *Nicotiana tabacum* cv Samsun using genetic constructs designed to direct the protein to the chloroplast IEM (Rolland *et al*., 2016). Two SbtA2 expression constructs with different promoters (*viz*. the tomato Rubisco small subunit promoter; *Sl*RbcS2, and the Arabidopsis ubiquitin promoter; *At*Ub10i) were generated to express an HAH6-tagged form of SbtA2-WH5701 (SbtA2-HAH6-WH5701) in *N. tabacum* cv Samsun.

Due to a lack of specific antibody for SbtA2, it was imperative to add an epitope tag to SbtA2-WH5701 in order to detect protein expression. Interestingly, adding a tag at the C-or N-terminus resulted in transporter non-function. However, we demonstrated that SbtA2-WH5701 carrying a tag located in the flexible cytoplasmic loop between transmembrane helices five and six (hereafter the 5-6 loop) had similar HCO_3_^-^ uptake activity as untagged SbtA2-WH5701 (Figure S3, S4). Using this epitope, we were able to determine the relative level of SbtA2-WH5701 expression in transgenic T_2_ *N. tabacum* (Tob^SbtA2^) plants. SbtA2-HAH6 protein expression in Tob^SbtA2^plants differed across their developmental age, dependent upon the promoter used. SbtA2 protein was more abundant in younger leaves of Tob^SbtA2^ plant lines using the *Sl*RbcS2 promoter, while those lines using the *At*ubi10i promoter had higher SbtA2 abundance in older leaves (Figure S6).

Tob^SbtA2^ plants appeared similar to wild type (Tob^WT^) with little to no bleaching of leaves throughout their lifespan (Figure 6H), which has been observed previously in *N. tabacum* expressing SbtA1 proteins (Figure S9). Compared to Tob^WT^, no differences were observed in relation to photosynthetic performance (CO_2_ assimilation rates and derived CO_2_ compensation point), final height, or whole plant fresh weight for Tob^SbtA2^ plants (Figure 6). While one *At*ubi10i line (Tob^SbtA_ubi-1.1.3^) demonstrated an elevated dry weight compared to Tob^WT^ (Figure 6C), and two Tob^SbtA2^ lines had significantly lower Δ^13^C values than Tob^WT^ (Figure 6G), these traits were not significantly different across all lines tested.

**Figure 6.**
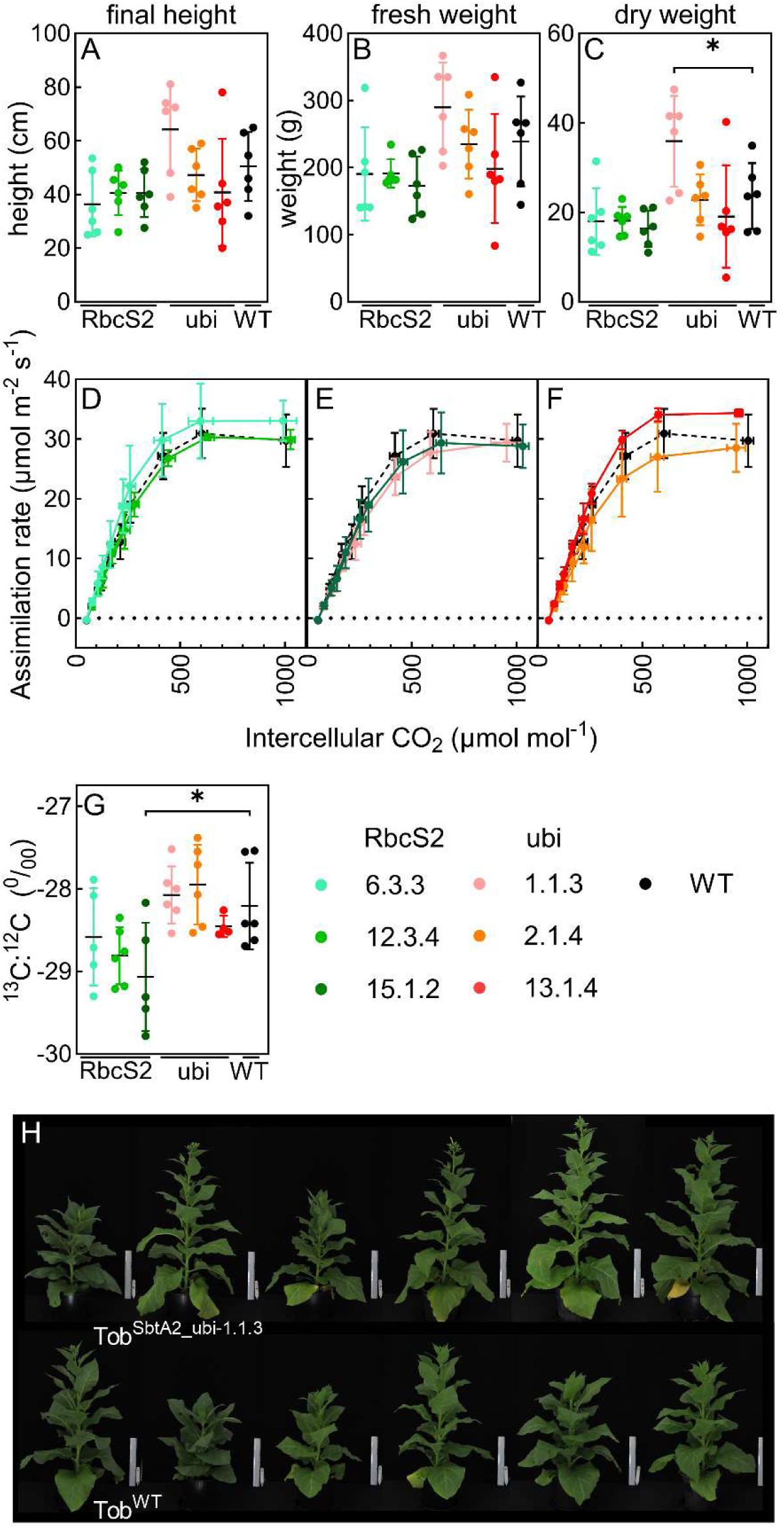
Phenotypic measurements of *N. tabacum* expressing SbtA-HAH6-WH5701 (Tob^SbtA^) compared to wild-type (Tob^WT^) plants. **(A)** The heights of plants were measured 63 days post cotyledon emergence (pce) from the base of the plants to the base of the apical meristem. The data is presented as means ± SD; n=6. **(B)** Each plant was harvested at the base of the stem 63 days pce and the fresh weight was measured. **(C)** The harvested plants were dried for 7 days at 80°C and the weight were measured. Data presented as mean ± SD; n=6. **(D-F)** A/Cint curves were measured on plants 30 ± 5 cm in height on young fully expanded leaves of similar size using a Licor 6800 with the following parameters; photon flux density 1500 μmol.m^-2.^s^-1^, at 25°C and photosynthetic measurements taken at 400, 200, 100, 50,150, 250, 400, 600, 800, 1200, 400 μmol.mol^-1^ CO2. Data presented as mean ± SD. n= 3-5 for Tob^SbtA2^ and n=9 for Tob^WT^. **(G)** Δ^13^C was determined from dry matter samples taken when the plants were 30 ± 5 cm in height. Asterisk (*) indicates significant difference between Tob^SbtA^ and Tob^WT^ plants as determined using unpaired *t*-test (P>0.05). Data presented as mean ± SD; n=4-6. **(H)** Comparison of Tob^SbtA-ubi^ and Tob^WT^ plants 63 days post cotyledon emergence.

## Discussion

The open-ocean α-cyanobacteria of the *Prochlorococcus* and *Synechococcus* (SynPro) clade is attributed with significant contributions to global CO_2_ fixation (Flombaum *et al*., 2013). Based on the enzyme kinetics of Form-IA Rubisco found in these organisms, cells require the accumulation of at least 10 mM HCO_3_^-^ to approach saturation of CO_2_ fixation (>200 µM CO_2_) within their α-carboxysomes (Whitehead *et al*., 2014; Shih *et al*., 2016). In the oceans, available CO_2_ levels are less than 15 µM (Young *et al*., 2016), although HCO_3_^-^ is stably available at around 2 mM (Falkowski and Raven, 2007). This highlights that HCO_3_^-^ uptake has evolved as a feature of biophysical CCMs in many oceanic single-celled algae and cyanobacteria, otherwise conditions would be severely limiting if dependent on diffusive CO_2_ uptake alone (Ritchie *et al*., 1996; Beardall and Raven, 2017). To date, most reports on cyanobacterial HCO_3_^-^ transporters have been on those originating from β-cyanobacteria, and relatively little is known about the Ci uptake systems that power the CCMs of the ecologically important α-cyanobacterial SynPro clade.

The type and number HCO_3_^-^ transporters differ in α- and β-cyanobacteria (Cabello-Yeves *et al*., 2022; Rae *et al*., 2011). Of the functional cyanobacterial HCO_3_^-^ transporters known (BCT1, SbtA1, BicA), generally the α-cyanobacterial *Synechococcus* clade has genes for one BicA HCO_3_^-^ transporter, while *Prochlorococcus* species have no genes for these HCO_3_^-^ transporters (Cabello-Yeves *et al*., 2022; Rae *et al*., 2011). Only distant homologues of SbtA1 and BicA, SbtA2 and BicA2, are apparent in SynPro (Rae *et al*., 2011). SbtA2 protein sequence identity is only approximately 25% compared to SbtA1 (Figure S1), however, its predicted structure (AlphaFold 3; Abramson *et al*., 2024) maps with high confidence to that of SbtA1 (Synechocystis PCC6803; PDB: 7egl; Fang *et al*., 2021 ; Figure S3A-C). This led us to hypothesise that SbtA2 likely functions as a HCO_3_^-^ transporter in species belonging to the SynPro clade.

### SbtA2 functional characteristics

In the present study we show for the first time that several members of the SbtA2 family from *Synechococcus* spp. are indeed functional HCO_3_^-^ transporters and displayed an ‘always-active’ HCO_3_^-^ transport function when expressed in *E. coli* (Figure 2). Compared to SbtA1-7942, SbtA2-WH5701 has a high HCO_3_^-^ transport flux (Figure 2B, 3A, 3C), in fact the highest we have recorded. In the presence of 200 mM NaCl, SbtA2-WH5701 had greater than six times the flux of SbtA1-7942 (Figure 3C). The maximum uptake flux by SbtA2 of 2.5 - 3 µmol.Ci OD_600_^-1.^h^-1^ generates an internal Ci pool size of up to 23.5 mM in *E. coli* over a 30 second period, and an accumulation factor of up to 70-fold compared to *E. coli* empty vector controls (Table 2). This extent of accumulation, if translated to a cyanobacterium or a C_3_ chloroplast, would be more than enough to reach the levels required to optimally operate the cyanobacterial CCM (Whitehead *et al*., 2014).

**Table 2:**
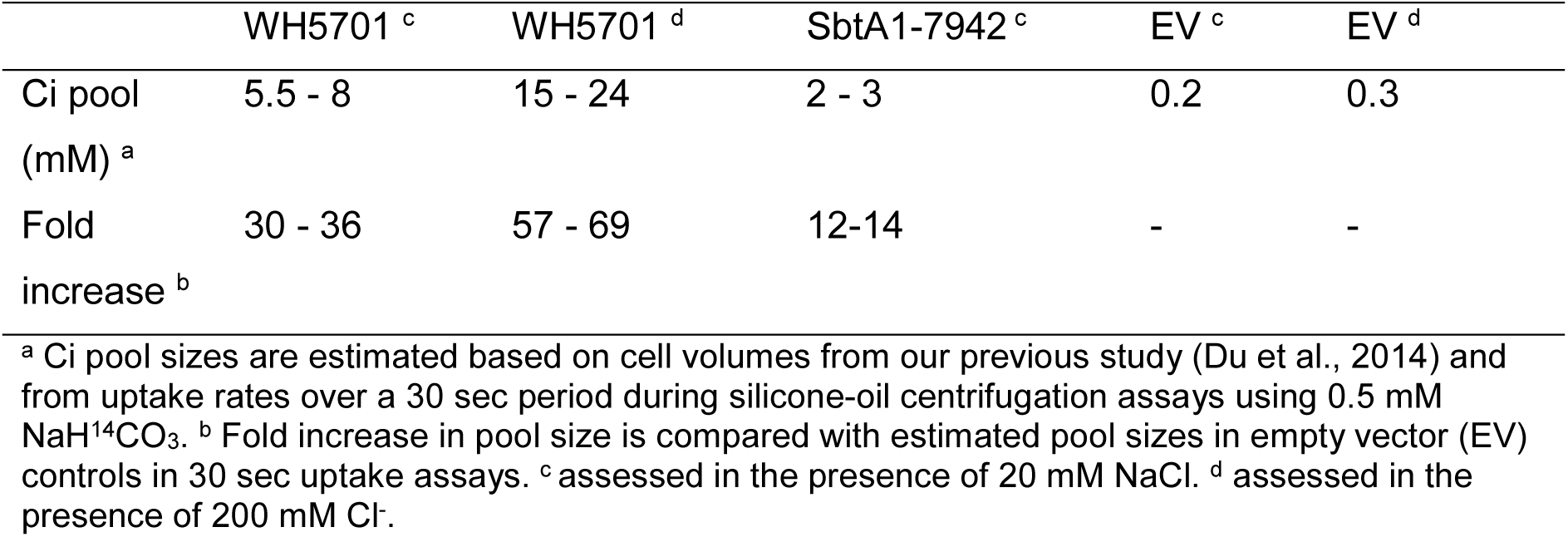
The estimated Ci pool sizes of *E. coli* cells expressing SbtA2 homologues.

|  | WH5701 <sup>c</sup> | WH5701 <sup>d</sup> | SbtA1-7942 <sup>c</sup> | EV <sup>c</sup> | EV <sup>d</sup> |
| --- | --- | --- | --- | --- | --- |
| Ci pool<br>(mM) <sup>a</sup> | 5.5 - 8 | 15 - 24 | 2 - 3 | 0.2 | 0.3 |
| Fold<br>increase <sup>b</sup> | 30 - 36 | 57 - 69 | 12-14 | - | - |

<sup>a</sup> Ci pool sizes are estimated based on cell volumes from our previous study (Du et al., 2014) and from uptake rates over a 30 sec period during silicone-oil centrifugation assays using 0.5 mM $\text{NaH}^{14}\text{CO}_3$ . <sup>b</sup> Fold increase in pool size is compared with estimated pool sizes in empty vector (EV) controls in 30 sec uptake assays. <sup>c</sup> assessed in the presence of 20 mM NaCl. <sup>d</sup> assessed in the presence of 200 mM $\text{Cl}^-$ .

Kinetic characterisation revealed that SbtA2 is a medium affinity transporter compared with those previously described (Rottet *et al*., 2021) exhibiting K_m_ values of approximately 150 - 200 µM (Table 1). When measured in *E. coli*, SbtA1 from *Cyanobium* PCC7001 and *Picosynechococcus* PCC7002 have affinities for HCO_3_^-^ of 189 µM and <100 µM respectively (Du *et al*., 2014). However, it is likely that more accurate affinities can be measured in model cyanobacteria in future studies since respiratory CO_2_ production in heterotrophic *E. coli* can lead to errors in affinity estimates (Du *et al*., 2014). For instance, the estimates of SbtA1 affinity for HCO_3_^-^ when expressed in cyanobacteria has been measured to be ∼2 µM (Price *et al*., 2004) which is much lower than when measured in *E. coli*. In comparison to the high affinity SbtA1 family, members of the low affinity transporter BicA, have K_m_ for HCO_3_^-^ estimated to be 75-353 µM when measured in cyanobacteria (BicA from *Synechococcus* WH8102, *Picosynechococcus* PCC7002 and *Synechocystis* PCC6803; Price *et al*., 2004).

Recognizing that the K_m_ of SbtA2 is well below the Ci concentration present in seawater (Falkowski and Raven, 2007), and the widespread nature of this gene family in α-cyanobacteria, leads us to believe that SbtA2 members could play a critical role in CCMs in this globally important cyanobacterial group. Furthermore, assuming sufficient expression, this indicates that SbtA2 could readily achieve the cytoplasmic HCO_3_^-^ accumulation known to be required in cyanobacteria for efficient CCM activity (Whitehead *et al*., 2014).

### Activation/Energisation of SbtA2 HCO_3_^-^ transport

From microbes to animals, HCO_3_^-^ transporters have been shown to function in a multitude of modes, including; HCO_3_^-^/SO_4_^2-^ and HCO_3_^-^/Cl^-^ exchangers (some of which are Na^+^-dependent; Cordat and Reithmeier, 2014), Na^+^/HCO_3_^-^ symporters, K^+^-dependent transporters (Parker and Boron, 2013), and ABC-type transporters that rely on ATP for HCO_3_^-^ transport (Omata *et al*., 1999; Gao *et al*., 2015). This highlights that, mechanistically, several drivers for HCO_3_^-^ uptake or exchange are entirely feasible and offer potential conceptual frameworks for SbtA2 function.

SbtA1 has a distinct requirement for Na^+^ for function, typically requiring 10 – 50 mM Na^+^ for maximum activity when expressed in *E. coli*, and likely a Na^+^/HCO_3_^-^ symporter (Shibata *et al*., 2002; Du *et al*., 2014). Despite SbtA2 having a predicted structure similar to SbtA1 (Figure S3A-C), evidence here suggests that Na^+^ does not drive SbtA2 activity. Instead, both Cl^-^ or SO_4_^2-^ might potentially drive SbtA2 function (Figure 3D, E), noting that Cl^-^ is substantially more abundant in seawater than SO_4_^2-^ (Nessim *et al*., 2015). SbtA2 demonstrated a high basal rate of HCO_3_^-^ transport when assayed in *E. coli*, which was not improved with additional Na^+^ (Figure 3). Moreover, the addition of 100 mM Cl^-^ or SO_4_^2-^ to the assay buffer (in the form of NaCl, KCl, LiCl, MgCl_2,_ MgSO_4_) resulted in an increase in the HCO_3_^-^ transport flux, approximately doubling the HCO_3_^-^ uptake activity compared to when no additional Cl^-^ or SO_4_^2-^ was added to the uptake buffer (Figure 3).

In the heterologous system used here, it is difficult to identify the exact mechanism enabling SbtA2 to transport HCO_3_^-^. Clearly further work in alterative systems is required to confirm the driving ion flux. Below we propose scenarios as to how Cl^-^ or SO_4_^2-^ may act on SbtA2 function.

### HCO_3_^-^/Cl^-^ antiport

The results presented in here are consistent with the possibility that SbtA2 may act to exchange HCO_3_^-^ with Cl^-^ or SO_4_^2-^, moving Cl^-^ or SO_4_^2-^ out of the cell as HCO_3_^-^ is moved in. There are multiple examples of HCO_3_^-^/Cl^-^ and HCO_3_^-^/SO_4_^2-^ exchangers (Zhang *et al*., 2023; Reithmeier, 2001; Reithmeier *et al*., 2016; Jennings, 2021). Evidence suggests that Cl^-^ can accumulate in *E. coli* during growth (Schultz *et al*., 1962), likely via H^+^/Cl^-^ exchangers that can actively transport Cl^-^ (eClC, Garcia-Celma *et al*., 2013), meaning that cells would have a Cl^-^ pool capable of driving HCO_3_^-^ uptake via HCO_3_^-^/Cl^-^ exchange. This could explain the high basal rate of HCO_3_^-^ transport of SbtA2 in *E. coli*, which is further stimulated with addition of Cl^-^ (Figure 3), if such an antiport mechanism were in operation. For this mode of action to be observed in *E. coli* there would need to be membrane transport systems enabling rapid import of Cl^-^ to increase HCO_3_^-^ uptake activity by SbtA2. Blocking these Cl^-^ transporters or measuring their net flux during HCO_3_^-^ uptake may provide evidence that this type of mechanism is in action. For such a mechanism to occur in oceanic cyanobacteria, however, cells would need to hyperaccumulate Cl^-^ from an external environment that already contains ∼500 – 600 mM Cl^-^ (Nessim *et al*., 2015) in order to enable gradient-driven HCO_3_^-^/Cl^-^ antiport. While Cl^-^ uptake has been observed (Ritchie, 1992), a mechanism to accumulate and maintain such high internal Cl^-^ would be required, and seems unlikely (Hagemann, 2011).

### HCO_3_^-^/Cl^-^ symport

In contrast, it may be possible that SbtA2 acts as a HCO_3_^-^/Cl^-^ symporter, transporting Cl^-^ and HCO_3_^-^ into the cell simultaneously. This seems less likely due to the resulting hyperpolarising effect of the ion influx on the membrane potential of the SbtA2-expressing organism, especially within a marine context. However, this may be possible if there are also active efflux mechanisms involved in the system to regulate cellular Cl^-^ concentration, preventing hyperpolarization and mitigating the ionic effects of SbtA2 function. Observed rates of HCO_3_^-^ uptake increased when exogenous Cl^-^ was experimentally elevated in *E. coli* expressing SbtA2 (Figure 3C-E), possibly supporting HCO_3_^-^/Cl^-^ symport as a potential mode of action. However, evidence against this mechanism is supported by the elevated base HCO_3_^-^ uptake rate of SbtA2 in the absence of SO_4_^2-^ or Cl^-^ (when Hepes buffered solution with pH adjusted to 7.5 with KOH was used as the assay buffer; Figure S7). In addition, existing literature does not clearly define HCO_3_^-^/Cl^-^ symport as a likely scenario.

### Allosteric control by Cl^-^/SO_4_^2-^

It is possible that allosteric control by Cl^-^/SO_4_^2-^ leads to SbtA2 activation. Evidence exists for this mechanism in alternate organisms (Davis-Kaplan *et al*., 1998; Tulloch *et al*., 2008; Perutz *et al*., 1994) and therefore sets a precedence for this type of control. In this scenario, the presence of Cl^-^/SO_4_^2-^ would alter protein conformation, oligomerisation, or charge around the active site to activate SbtA2, thus enabling transport function. Since SbtA2 exhibits Michaelis-Menten kinetics in Ci response assays (Figure 4 and S4), it is unlikely that SbtA2 is a passive channel and that after allosteric activation by Cl^-^ a secondary counter-ion (K^+^, Na^+^, Mg^2+^) must be required for HCO_3_^-^ transport by SbtA2. If this is the mode of activation for SbtA2, the site of Cl^-^/SO_4_^2-^ elevation (intracellular or extracellular), the mechanism of Cl^-^/SO_4_^2-^ action on SbtA2, and the secondary counter-ion would need to be determined to optimise function in non-native environments.

These speculative ideas need further investigation to clarify the mode of action of SbtA2, perhaps using ^36^Cl^-^ to assess Cl^-^ flux in *E. coli*, or by expressing SbtA2 in *Xenopus* oocytes, or in liposomes where other membrane transport systems may be less likely to confound results. Without further analysis we cannot yet conclude what is the energisation mode for SbtA2. Additionally, expression of SbtA2 in the *Δ5* mutant of *Synechocystis* sp. PCC6803, which has all five Ci uptake mechanisms deleted (Xu *et al*., 2008), may provide functional information in a cyanobacterial context.

### Prochlorococcus SbtA2

Curiously, the two SbtA2 members from *Prochlorococcus* (Figure 2A) did not function in *E. coli* despite the high sequence similarity of all five homologues (Figure S1). There may be three things to consider here. Firstly, SbtA2’s from *Prochlorococcus* may be pseudo-genes, though this seems unlikely since there are so few genes coding for putative HCO_3_^-^ transporters in *Prochlorococcus* genomes (Rae *et al*., 2011). Secondly, there may be another level of allosteric control that cannot be satisfied in the heterologous *E. coli* expression system. If so, this will require further work to resolve the functional activation of SbtA2 from *Prochlorococcus*. Using an approach such as laboratory evolution to generate functional HCO_3_^-^ transporters has been shown to be useful to resolve important sequence elements contributing to both HCO_3_^-^ transporter and CA function (Pulsford *et al*., 2024; Rottet *et al*., 2024). This was attempted on the non-functional members; however, no functional forms have been generated to date in our hands. Thirdly, *Prochlorococcus* species appear to use sulpholipids bilayers (Van Mooy *et al*., 2006) rather than phospholipid bilayers found in *E. coli*, so there is uncertainty whether *Prochlorococcus* membrane proteins will insert or interact correctly within bacterial membranes. The importance of *Prochlorococcus* as an oceanic contributor to carbon acquisition, and the lack of evidence of any HCO_3_^-^ transport in this genus is intriguing and led us to question how these organisms can be achieving efficient CO_2_ fixation in low CO_2_ environments without possessing any characterised HCO_3_^-^ transporters. On balance, the most likely scenario is that SbtA2, and/or the distant BicA homologue, are functional HCO_3_^-^ transporters in *Prochlorococcus* spp. or there are alternative HCO_3_^-^ transporters or regulators present that are still undiscovered.

### SbtA2:B2 interaction

The SbtB1 protein has been shown to regulate the SbtA1 transporter as a “curfew protein” that inhibits the action of the pump (Förster *et al*., 2023; Kaczmarski *et al*., 2019; Fang *et al*., 2021). Additionally, SbtB1 appears to act as a nucleotide- and redox-sensitive regulator that controls SbtA1 by switching between binding states and promoting AMP-dependent complex formation, possibly modulating HCO_3_^-^ transport in response to cellular energy and light/dark conditions (Selim *et al*., 2023). In either case, the system is regulated by cellular ATP, or the relative adenylate charge, such that high ATP concentrations lead to the unbinding of the SbtB1 trimer from the SbtA1 transport trimer, thereby activating SbtA1 (Förster *et al*., 2023). Previously, Du *et al*. (2014) showed that SbtA1 activity could be eliminated through co-expression of SbtB1 in *E. coli* (DH5α and EDM636 strains). This implies that ATP concentrations inside growing *E. coli* cells must be low enough for SbtB1 to bind SbtA1, resulting in the prevention of SbtA1 function and no growth of *E. coli* (EDCM636 strain) at atmospheric CO_2_. Here we found that SbtB2 can inhibit its cognate SbtA2 function by around 30-90% (Figure 5 and S5), depending on protein expression levels, and that the expression level of SbtB2 is important in controlling SbtA2 function. Co-expression of SbtA2:B2 in CAfree did not prevent growth of CAfree *E. coli* at atmospheric CO_2_ (Figure 5A and S2B). However, HCO_3_^-^ uptake assays in *E. coli* DH5α demonstrated reduced levels of uptake (Figure 5, and S5), and highlights that this is likely sufficient to overcome the CAfree phenotype in growth studies.

### The effect of epitope tagging on SbtA2 function

Efforts to generate antibodies to SbtA2 for use in immunoblot identification of heterologously expressed protein was unsuccessful. We initially attempted to introduce epitope tags to the N- or C-terminus of SbtA2 but found these resulted in non-functional transporters (Figure S8). Using the predicted structure of SbtA2 (Abramson *et al*., 2024) and the crystal structure of SbtA1 (Fang *et al*., 2021; Liu *et al*., 2021) as guides (Figure S3), we were able to conclude that the cytoplasmic 5-6 loop is a disordered flexible region that may be amenable to epitope tagging. Functional data from tagging studies of SbtA1-7942 suggested this region was not functionally critical in that form of SbtA1 (Du *et al*., 2014). Here, all assessments of SbtA2 forms carrying an internal epitope-tag within this flexible cytoplasmic 5-6 loop region indicated that these proteins retained function comparable to their untagged counterparts (Figure S3). This understanding was critical to using these SbtA2 forms to assess protein expression in both *E. coli* and tobacco without loss of HCO_3_^-^ transport activity.

### Expression of SbtA2 in *N. tabacum*

Introduction of CCMs to chloroplasts of C_3_ plants has been modelled to boost Rubisco carboxylation, increasing photosynthetic efficiency, thereby improving crop productivity (Price *et al*., 2011; Rottet *et al*., 2021; McGrath and Long, 2014) and a possible strategy to circumvent the looming food shortages expected as global population increases (Ray *et al*., 2013). The expression of functional HCO_3_^-^ transporters in the chloroplast IEM could support a 7-16% increase in crop productivity resulting from increased CO_2_ supply to Rubisco (Price *et al*., 2011; McGrath and Long, 2014). Previously several HCO_3_^-^ transporters have been expressed in the chloroplast IEM of model plants, although with mixed results with respect to improvement in photosynthetic parameters. For the most part, there has been no evidence of active chloroplast HCO_3_^-^ accumulation, and where there have been improvements in photosynthetic parameters, this has not been clearly linked to increased CO_2_ supply to Rubisco (Rottet *et al*., 2024; Forster *et al*., 2023; Nölke *et al*., 2019; Atkinson *et al*., 2016; Pengelly *et al*., 2014).

Here we successfully directed the epitope-tagged SbtA2-HAH6-WH5701 to the chloroplast IEM of *N. tabacum* after nuclear expression (Figure S6). T_2_ plants were assessed for phenotypic and photosynthetic changes to determine the impact of the expressed SbtA2 protein. Although one Tob^SbtA2^ expression line had increased dry matter and two with lower Δ^13^C (Figure 6G), there was no consistent improvement of photosynthetic parameters or phenotypic traits of plants expressing SbtA2 compared to wild type (Figure 6). Notably, however, transgenic plant phenotypes appeared similar to wild type plants, and no detrimental effects on plant growth was observed. This contrasts with previous attempts to express SbtA1 in *N. tabacum* which were slow growing and demonstrated a chlorotic phenotype (Figure S9). We conclude from this that SbtA2 may either be non-functional *in planta*, or that the level of function is too low to provide any observable benefit in the lines analysed. It is possible that an alternate SbtA2 homologue may be a better candidate for expression *in planta* (for example SbtA2-CB0205 which appears to have higher activity per unit protein than SbtA2-WH5701 [Figure S3E, F]), or that plant lines showing higher expression across all stages of plant growth might provide a better indication of expected photosynthetic improvements.

It still needs to be established with greater certainty if SbtA2 members are in fact Cl^-^/HCO_3_^-^ (and/or SO_4_^2-^/HCO_3_^-^) antiporters, or indeed if they are under Cl^-^/SO4^-^ allosteric control. If SbtA2 does rely on Cl^-^ (or SO_4_^2-^) for HCO_3_^-^ transport function, it is important to understand the existing requirement, magnitude, and maintenance of a Cl^-^ (or SO_4_^2-^) gradient between the cytoplasm and the chloroplast stroma. This also highlights the pressing need to understand the Cl^-^ and SO_4_^2-^ concentration gradients and movement in leaf tissue sub-compartments and the endogenous transporters involved. This will determine whether the appropriate activation or energisation conditions exist for SbtA2 function in the chloroplast IEM.

It is understood that Cl^-^ is the most abundant anion in the chloroplast (Colmenero-Flores *et al*., 2019) and serves an important role in photosynthesis as an essential cofactor in photosystem II function (Colmenero-Flores *et al*., 2019). Cl^-^ concentrations have been measured at 36-60 mM in chloroplasts of the glaucophyte spinach (Demmig and Gimmler, 1983). Interestingly, similar Cl^-^ concentrations have been measured in chloroplasts of other important C_3_ plants, barley and wheat (39 and 48 mM Cl^-^ respectively; Robinson and Downton, 1984). Additionally, tobacco can accumulate Cl^-^ in macronutrient quantities without apparent detrimental effects on the plants when grown in elevated Cl^-^ as long as particular cations are not in excess (Franco-Navarro *et al*., 2019). Furthermore, growth in elevated NaCl can increase chloroplastic Cl^-^ concentrations in halophytic plants (Demmig and Winter, 1986). This is an example of a factor that could be optimised in future plant expression studies, if SbtA2 is indeed a Cl^-^/HCO_3_^-^ antiporter, or in some way linked to Cl^-^ availability. However, there are a number of other issues that need optimisation, such as; achieving sustained expression and insertion of SbtA2 targeted to the chloroplast envelope inner membrane, studies to ascertain if a sufficient electrochemical gradient exists between the cytosol and chloroplast stroma, and whether this can be optimised by expression of CLC-type Cl^-^ importers.

## Conclusion

This study involved characterisation of a previously undescribed family of HCO_3_^-^ transporters, SbtA2, chiefly found in α-cyanobacteria (Figure 1). Although SbtA2 appears to be a distant relative of the Na^+^-dependent SbtA1, the mode of action of SbtA2 is quite different, with an apparent dependency on Cl^-^ or SO_4_^2-^. SbtA2 also displays high HCO_3_^-^ transport rates, and an affinity for HCO_3_^-^ that can be described as intermediate with respect to other known HCO_3_^-^ transporters. The functional characterisation of SbtA2 in *E. coli* presents as an ‘always active’ phenotype, which lends itself as a potential candidate for insertion into the IEM of the chloroplast with the aim of improving photosynthetic efficiency in C_3_ crop plants. However, *N. tabacum* expressing SbtA2 showed no consistent improvement in photosynthetic parameters compared to wild type plants. Nonetheless, this work highlights that a broader selection of HCO_3_^-^ transporters could have the potential to enhance photosynthetic performance in C_3_ plants by delivering Ci to the chloroplast. In order for successful function of SbtA2 *in planta* it is important to understand the apparent Cl^-^/SO_4_^2-^ dependency of SbtA2 and the concentrations of these ions in sub-compartments in plant leaf tissue, and the consequences of over-expressing SbtA2 and any related ion imbalances this may cause.

## Abbreviations

CA: carbonic anhydrase
CAfree: pecialized *E. coli* strain that lacks CAs
CCM: CO_2_-concentrating mechanism
CO_2_: carbon dioxide
Ci: inorganic carbon
C_int_: intercellular CO_2_
HCO_3_^-^: bicarbonate
IEM: inner envelope membrane
IPTG: isopropyl β-D-1-thiogalactopyranoside LB Luria-Bertani
SbtA: sodium-dependent bicarbonate transporter

## Supplementary Data

The following supplementary data are available online. Supplementary Method Epitope tagging of SbtA2.

Table S1 Addition of epitope tags to the 5-6 cytoplasmic loop of SbtA2. Table S2 Affinity of H6-tagged SbtA2 homologues for HCO_3_^-^.

Table S3 Antibodies used in this study.

Figure S1 Alignment of SbtA homologues across cyanobacteria.

Figure S2 Liquid growth assay of CAfree expressing SbtA2±B2.

Figure S3 Internal epitope tagging of SbtA2 and functional analysis.

Figure S4 HCO_3_^-^ uptake kinetics of tagged and untagged forms of SbtA2.

Figure S5 SbtAB2 gene arrangements and SbtB2 co-expression effects on SbtA2 HCO_3_^-^ uptake in *E. coli*.

Figure S6 Assessment of the localisation and expression of SbtA2 in *N. tabacum*.

Figure S7 Analysis of the basal HCO_3_^-^ uptake rate of SbtA2

Figure S8 Assessment of N- and C-terminally tagged SbtA2 HCO_3_^-^ transport function and protein expression.

## Acknowledgements

The authors thank David Savage (UC Berkeley) for kindly providing the CAfree strain of *E. coli.* We also thank Suyan Yee and Sarah Rottet for carrying out confocal and transient expression analysis, and Kelly Chapman for technical assistance with the Licor6800.

## Author contributions

LMR, GDP, BML: conceptualization and visualisation; GDP project administration; LMR: methodology and analysis; BML, GDP, CSB: supervision; LMR: writing – original draft; all authors: writing – review and editing; GDP: funding acquisition.

## Conflict of interest

The authors declare no conflicts of interest.

## Funding Statement

LMR received an Australian Government Research Training Program Scholarship to cover tuition fees.

## Data availability

The primary data supporting this study are publicly available at doi: 10.17632/wrwjkyn6bn.1 (Rourke *et al*., 2025) and are available from the corresponding author upon request. Additional data reported in this paper are presented in the Supplementary Data.

## Notes

### Competing Interest Statement

The authors have declared no competing interest.

### Summary of Updates

Addition of Supplementary data files.

