## Supplementary data for "Functional characterisation of bicarbonate transporters from the cyanobacterial SbtA2 family and subsequent expression in tobacco"

#### Supplementary Method: Epitope tagging of SbtA2

It was necessary to introduce epitope tags to the proteins for SbtA2:B2 binding studies and protein abundance determination when expressed in *E. coli* and *in planta*. A candidate location for epitope tag placement was identified within the 5-6 cytoplasmic loop (referred to herein as the '5-6 loop') by modelling SbtA2 against the structure of SbtA1 from *Synechocystis* PCC6803 (SbtA1-6803; Fang *et al.*, 2021; Figure S3A-C), and using known functional data from tagging studies of SbtA1 from *Synechococcus* PCC7942 (SbtA1-7942) that suggested this region was not functionally critical in the PCC7942 form of SbtA1 (Du *et al.*, 2014). We hypothesised that this flexible cytoplasmic loop may be amenable to additional sequence insertion without adversely affecting $HCO_3^-$ uptake function. Flexible glycine linkers flanking either His×6 (H6) or HAHis×6 (HAH6) epitope tags were added to this cytoplasmic loop, after amino acid A157 (referred to as SbtA2-H6 and SbtA2-HAH6 respectively; Figure S3 and Table S3).

#### The effect of tagging SbtA2 on $HCO_3^-$ uptake

The impact of epitope tags on SbtA2 $HCO_3^-$ transport function was assessed by comparing the $HCO_3^-$ uptake rates of tagged and untagged SbtA2. SbtA2-H6-WH5701 and SbtA2-HAH6-WH5701 had similar $HCO_3^-$ uptake activity to untagged SbtA2-WH5701 (Figure S3D). SbtA2-RCC307 and SbtA2-CB0205 were also tagged with H6 flanked by flexible 3×Gly linkers (SbtA2-H6-RCC307 and SbtA2-H6-CB0205 respectively) inserted after amino acid residues Arg156 and Asn157 respectively. SbtA2-H6-RCC307 and SbtA2-H6-CB0205 were similarly active to their untagged forms (Figure S3E).

This retention of activity by internally tagged SbtA2-H6 forms enabled the comparison of SbtA2 transporter activity with estimates of relative protein abundance of each SbtA2 homologue. Based on culture density, SbtA2-H6-WH5701 had the highest $HCO_3^-$ uptake activity (Figure S3E), however, SbtA2-H6-WH5701 protein was expressed in higher abundance than SbtA2-H6-RCC307, and more so than SbtA2-H6-CB0205 (Figure S3E, F). Assuming all protein detected was functional, we can infer that SbtA2-CB0205 is likely the highest flux transporter of the three SbtA2 members tested when $HCO_3^-$ uptake activity is based on protein expression.

The activity of the 5-6 loop-tagged SbtA2:B2 construct (SbtA2-H6:SbtB2-HA) was similar to the untagged SbtA2:B2-HA form (Figure S5E), this implied the tags on SbtA2 and SbtB2 did not interfere with SbtA2:B2 binding, which indicated SbtA2-H6:B2-HA would be suitable to assess SbtA2:B2 binding. Although the $HCO_3^-$ uptake experiments strongly indicate their association, *in vitro* binding studies provided minimal evidence of SbtA2-H6:B2-HA binding (Figure S5H)

### Supplementary Tables

Table S3: Addition of epitope tags to the 5-6 cytoplasmic loop of SbtA2.

| Tag | Epitope | Sequence | Function (% of WT) |
|---|---|---|---|
| WT | None | ASSAAA$^{157}$EQPR | 100% |
| H6 | G3-H6-G3 | A$^{157}$GGGHHHHHHGGGE | 87-110% |
| HAH6 | G3-HA-H6-G3 | A$^{157}$GGGYPYDVPDYAHHHHHHGGGE | 78-92% |

The $HCO_3^-$ uptake activity of DH5α strain of E. coli expressing SbtA2-WH5701 (WT), SbtA2-H6-WH5701 (H6) and SbtA2-HAH6 (HAH6) were assessed. The activity of the tagged forms are shown relative to the untagged SbtA2-WH5701, n=12 from 3 biological replicates. Uptake assay buffer contained 20 mM BTP-$H_2SO_4$ pH 7.5 + 20 mM NaCl.

Table S4: Affinity of H6-tagged SbtA2 homologues for $HCO_3^-$.

| | WH5701-H6 | RCC307-H6 | CB0205-H6 |
|---|---|---|---|
| $V_{max}$ | 1.39 ± 0.18 | 0.61 ± 0.08 | 0.36 ± 0.02 |
| $K_m$ (μM) | 155 ± 65 | 154 ± 10 | 157 ± 29 |

The $K_m$ (μM) and $V_{max}$ (μmol Ci·$OD_{600}^{-1}$·$h^{-1}$) of H6-tagged SbtA2 transporters in the presence of 20 mM NaCl in the assay buffer, mean from 3 assays.

Table S5: Antibodies used in this study.

| Antibody | Primary or secondary | Epitope sequence | Source | Dilution |
|---|---|---|---|---|
| His×6 | primary | HHHHHH | Cat # AS11 1771 Agrisera | 1:3000 |
| HA | primary | YPYDVPDYA | Cat # H9658 Merck Life Science, USA | 1:10000 |
| Flag | primary | DYKDDDDK | Cat # F1804 Merck Life Science, USA | 1:2000 |
| Goat-anti-mouse AP-conjugated | secondary | | Cat # A3562 Merck Life Science, USA | 1:3000 |

**Figures**

A

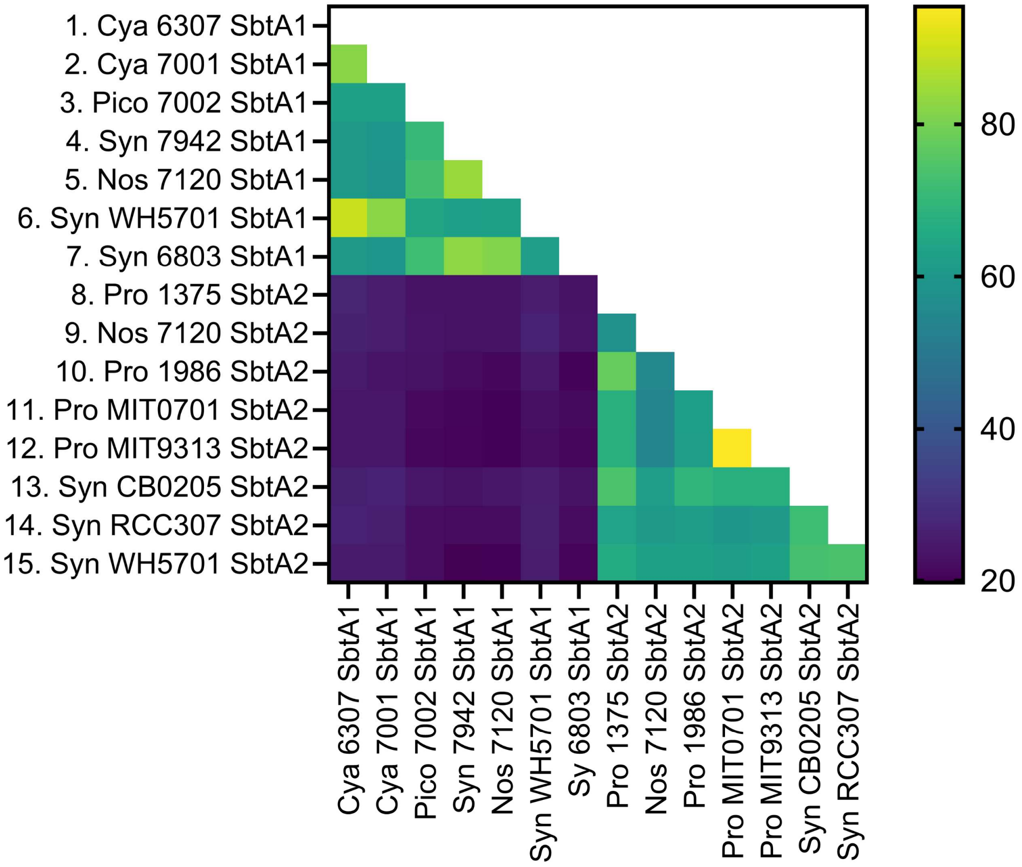

1. Cya 6307 SbtA1
2. Cya 7001 SbtA1
3. Pico 7002 SbtA1
4. Syn 7942 SbtA1
5. Nos 7120 SbtA1
6. Syn WH5701 SbtA1
7. Syn 6803 SbtA1
8. Pro 1375 SbtA2
9. Nos 7120 SbtA2
10. Pro 1986 SbtA2
11. Pro MIT0701 SbtA2
12. Pro MIT9313 SbtA2
13. Syn CB0205 SbtA2
14. Syn RCC307 SbtA2
15. Syn WH5701 SbtA2
Cya 6307 SbtA1
Cya 7001 SbtA1
Pico 7002 SbtA1
Syn 7942 SbtA1
Nos 7120 SbtA1
Syn WH5701 SbtA1
Sy 6803 SbtA1
Pro 1375 SbtA2
Nos 7120 SbtA2
Pro 1986 SbtA2
Pro MIT0701 SbtA2
Pro MIT9313 SbtA2
Syn CB0205 SbtA2
Syn RCC307 SbtA2
80
60
40
20

B

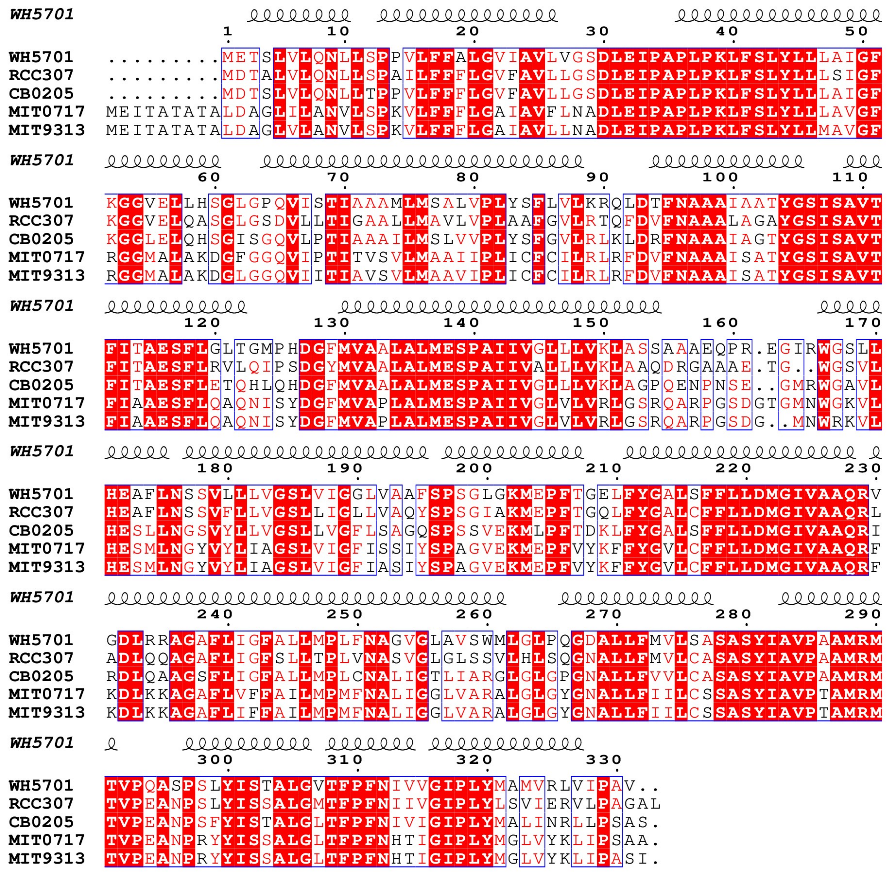
WH5701
WH5701 .........METSLVLQNLLSPPVLFFALGVIAVLVGSDLEIPAPLPKLFSLYLLLAIGF
RCC307 .........MDTALVLQNLLSPAILFFFLGVFAVLLGSDLEIPAPLPKLFSLYLLLSIGF
CB0205 .........MDTSLVLQNLLTPPVLFFFLGVFAVLLGSDLEIPAPLPKLFSLYLLLAIGF
MIT0717 MEITATATALDAGLILANVLSPKVLFFFLGAIAVFLNADLEIPAPLPKLFSLYLLLAVGF
MIT9313 MEITATATALDAGLVLANVLSPKVLFFFLGAIAVLLNADLEIPAPLPKLFSLYLLMAVGF
WH5701 KGGVELLHSGLGPQVISTIAAAMLMSALVPLYSFLVLKRQLDTFNAAAIAATYGSISAVT
RCC307 KGGVELQASGLGSDVLLTIGAALLMAVLVPLAAFGVLRTQFDVFNAAALAGAYGSISAVT
CB0205 KGGLELQHSGISGQVLPTIAAAILMSLVVPLYSFGVLRLKLDRFNAAAIAGTYGSISAVT
MIT0717 RGGMALAKDGFGGQVIPTITVSVLMAAIIPLICFCILRLRFDVFNAAAISATYGSISAVT
MIT9313 RGGMALAKDGLGGQVIITIAVSVLMAAVIPLICFCILRLRFDVFNAAAISATYGSISAVT
WH5701 FITAESFLGLTGMPHDGFMVAAALALMESPAIIVGLLLVKLASSAAAEQPR.EGIRWGSLL
RCC307 FITAESFLRVLQIPSDGYMVAALALMESPAIIVALLLVKLAAQDRGAAAE.TG..WGSVL
CB0205 FITAESFLETQHLQHDGFMVAALALMESPAIIVGLLLVKLAGPQENPNSE..GMRWGAVL
MIT0717 FIAAESFLQAQNISYDGFMVAPLALMESPAIIVGLVLVRLGSRQARPGSDGTGMNWGKVL
MIT9313 FIAAESFLQAQNISYDGFMVAPLALMESPAIIVGLVLVRLGSRQARPGSDG..MNWRKVL
WH5701 HEAFLNSSVLLLVGSLVIGGLVAAFSPSGLGKMEPFTGELFYGALSFFLLDMGIVAAQRV
RCC307 HEAFLNSSVFLLVGSLLIGLLVAQYSPSGIAKMEPFTGQLFYGALCFFLLDMGIVAAQRL
CB0205 HESLLNGSVYLLVGSLLVGFLSAGQSPSSVEKMLPFTDKLFYGALSFFLLDMGIVAAQRI
MIT0717 HESMLNGYVYLIAGSLVIGFISSIYSPAGVEKMEPFVYKFFYGVLCFFLLDMGIVAAQRF
MIT9313 HESMLNGYVYLIAGSLVIGFIASIYSPAGVEKMEPFVYKFFYGVLCFFLLDMGIVAAQRF
WH5701 GDLRRAGAFLIGFALLMPLFNAGVGLAVSWMLGLPQGDALLFMVLSASASYIAVPAAMRM
RCC307 ADLQQAGAFLIGFSLLTPLVNASVGLGLSSVLHLSQGNALLFMVLCASASYIAVPAAMRM
CB0205 RDLQAAGSFLIGFALLMPLCNALIGTLIARGLGLGPGNALLFVVLCASASYIAVPAAMRM
MIT0717 KDLKKAGAFLVFFAILMPMFNALIGGLVARALGLGYGNALLFIILCSSASYIAVPTAMRM
MIT9313 KDLKKAGAFLIFFAILMPMFNALIGGLVARALGLGYGNALLFIILCSSASYIAVPTAMRM
WH5701 TVPQASPSLYISTALGVTFPFNIVVGIPLYMAMVRLVIPAV..
RCC307 TVPEANPSLYISSALGMTFPFNIIVGIPLYLSVIERVLPAGAL
CB0205 TVPEANPSFYISTALGLTFPFNIVIGIPLYMALINRLLPSAS.
MIT0717 TVPEANPRYYISSALGLTFPFNHTIGIPLYMGLVYKLIPSAA.
MIT9313 TVPEANPRYYISSALGLTFPFNHTIGIPLYMGLVYKLIPASI.

C

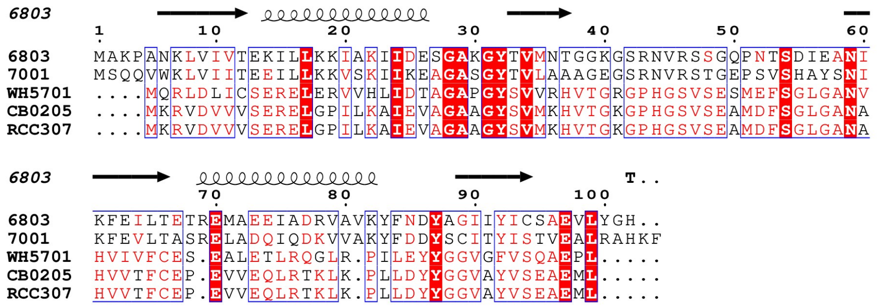
6803
6803 MAKPANKLVIVTEKILLKKIAKIIDESGAKGYTVMNTGGKGSRNVRSSGQPNTSDIEANI
7001 MSQQVWKLVIITEEILLKKVSKIIKEAGASGYTVLAAAGEGSRNVRSTGEPSVSHAYSNI
WH5701 ....MQRLDLICSERELERVVHLIDTAGAPGYSVVRHVTGRGPHGSVSESMEFSGLGANV
CB0205 ....MKRVDVVVSERELGPILKAIEVAGAAGYSVMKHVTGKGPHGSVSEAMDFSGLGANA
RCC307 ....MKRVDVVVSERELGPILKAIEVAGAAGYSVMKHVTGKGPHGSVSEAMDFSGLGANA
T..
6803 KFEILTETREMAEEIADRVAVKYFNDYAGIIYICSAEVLYGH..
7001 KFEVLTASRELADQIQDKVVAKYFDDYSCITYISTVEALRAHKF
WH5701 HVIVFCES.EALETLRQGLR.PILEYYGGVGFVSQAEPL.....
CB0205 HVVTFCEP.EVVEQLRTKLK.PLLDYYGGVAYVSEAEML.....
RCC307 HVVTFCEP.EVVEQLRTKLK.PLLDYYGGVAYVSEAEML.....

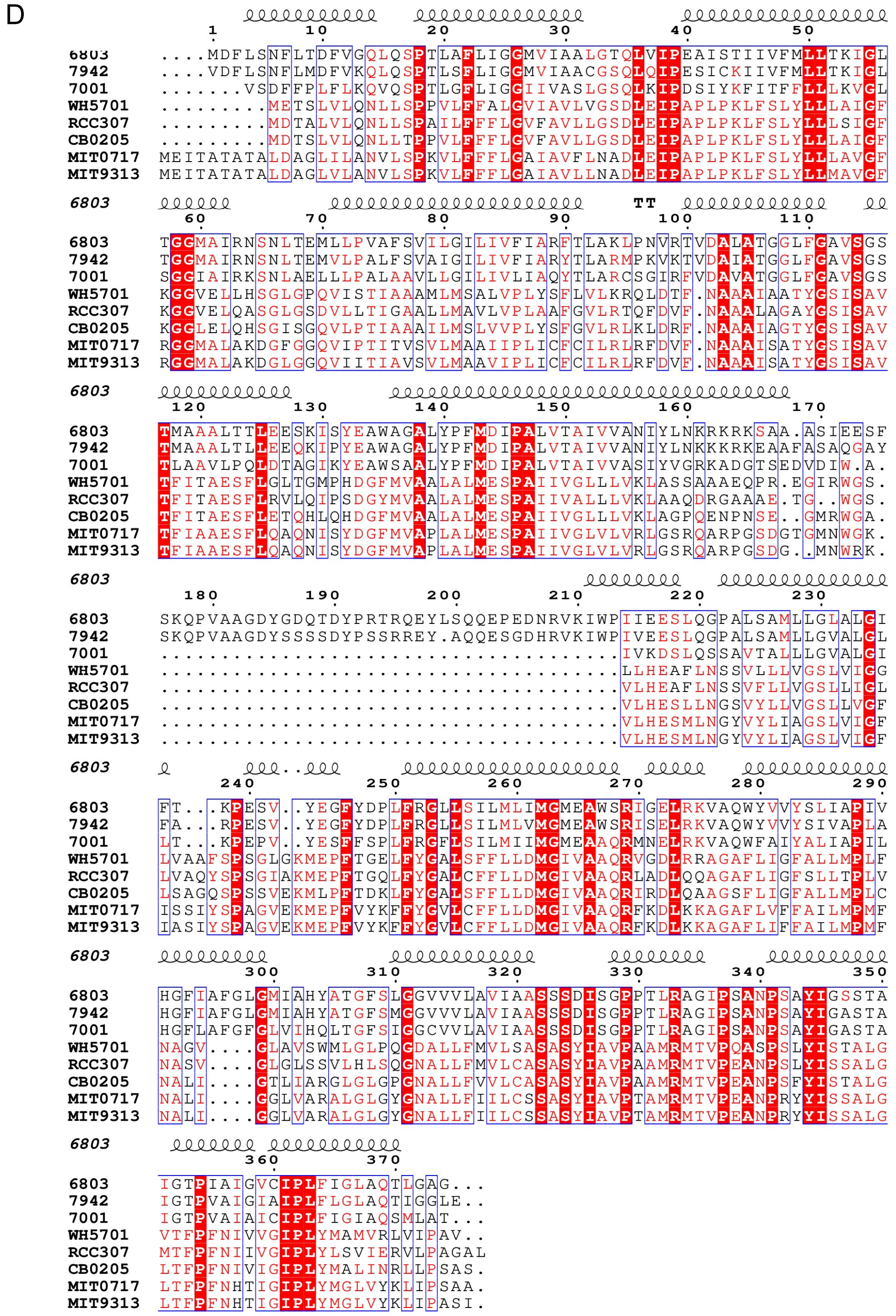


**Figure S1**: **Alignment of SbtA homologues across cyanobacteria**

**(A)** SbtA1 and SbtA2 have ~20-27% protein sequence identity. Species abbreviations: Pico, *Picosynechococcus*; Pro, *Prochlorococcus*; Syn, *Synechococcus*; Sy, *Synechocystis*; Cy, *Cyanobium*; Nos, *Nostoc*. Alignments performed by Geneious Prime software. **(B)** Alignment of SbtA2 from *Synechococcus* WH5701, CB0205, RCC307 and *Prochlorococcus* MIT0701 and MIT9313. **(C)** Alignment of SbtB1 from *Synechocystis* PCC6803, *Cyanobium* PCC7001 and SbtB2 from *Synechococcus* WH5701, CB0205, RCC307. **(D)** Alignment of SbtA1 from *Synechocystis* PCC6803, *Cyanobium* PCC7001 and *Synechococcus* PCC7942, and SbtA2 from *Synechococcus* WH5701, CB0205, RCC307 and *Prochlorococcus* MIT0701 and MIT9313. Alignments performed using Multalin (Corpet, 1988) and displayed using ESPript 3.0 (Robert and Gouet, 2014).

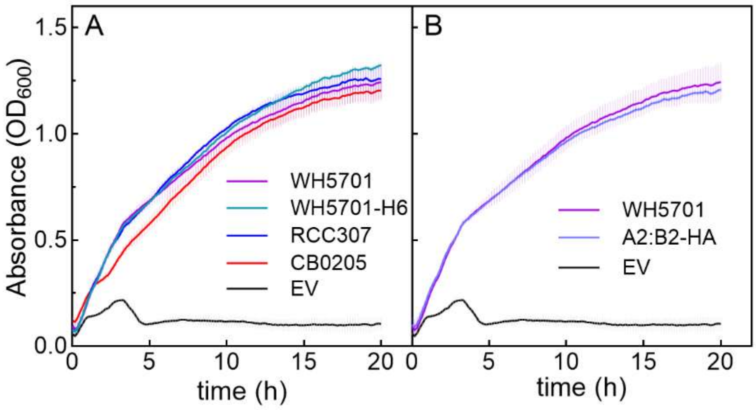


**Figure S2**: **Liquid growth assay of CAfree expressing SbtA2±B2**
**(A)** CAfree expressing SbtA2-WH5701 (WH5701), SbtA2-RCC307 (RCC307), SbtA2-CB0205 (CB0205) and a negative control (empty vector; EV) were grown in LB in ambient $CO_2$ for 20 h. Optical density ($OD_{600}$) was measured every 10 minutes. **(B)** CAfree expressing SbtA2-WH5701 (WH5701), SbtA2:B2-HA-WH5701 (A2:B2-HA) and a negative control (empty vector; EV) were grown in LB in ambient $CO_2$ for 20 h. Optical density ($OD_{600}$) was measured every 10 minutes.

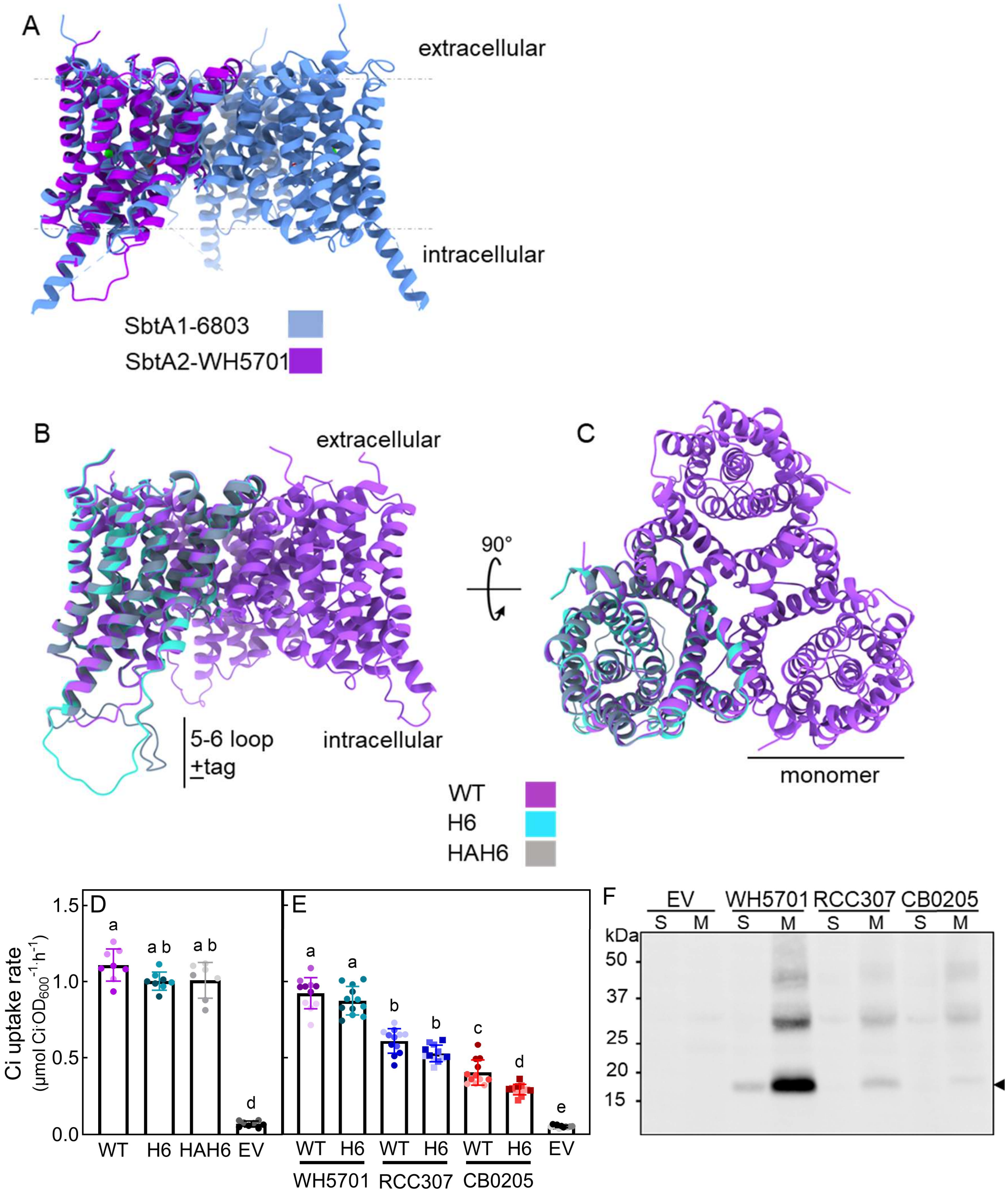


**Figure S3: Internal epitope tagging of SbtA2 and functional analysis.**
**(A)** The predicted structure of SbtA2-WH5701 was overlaid on to SbtA1 from *Synechocystis* PCC6803 (SbtA1-6803, PDB:7egl; Fang *et al.*, 2021). **(B-C)** The predicted structures of SbtA2-WH5701 monomers tagged with H6 (H6) and HAH6 (HAH6) in the 5-6 loop and a trimer of SbtA2-WH5701 (WT) were overlaid to demonstrate structural comparison of the proteins. The structures are shown as a membrane cross-section view in (B) and shown extracellularly in (C). SbtA2 structures were predicted using AlphaFold 3 (Abramson *et al.*, 2024) and visualised using ChimeraX (Pettersen *et al.*, 2021). **(D)** $HCO_3^-$ uptake activity of SbtA2-WH5701 (WT), SbtA2-H6-WH5701 (H6) and SbtA2-HAH6-WH5701 (HAH6) and a negative control (empty vector; EV). Assay buffer composition: 20 mM BTP-$H_2SO_4$ pH 7.5 + 20 mM NaCl. n=7-8 from 2 biological replicates. **(E)** $HCO_3^-$ uptake activity of untagged and tagged forms of SbtA2 from *Synechococcus* WH5701 (WT vs H6 tagged), RCC307 (WT vs H6 tagged), CB0205 (WT vs H6 tagged) and a negative control (empty vector; EV). SbtA2-WH5701 (A2). Assay buffer composition: 20 mM BTP-HCl pH 7.5. n=12 from 3 biological

replicates. **(F)** Immunoblot analysis of crude membrane-enriched (M) and soluble (S) fractions prepared from *E. coli* expressing SbtA2-H6 from *Synechococcus* WH5701, RCC307 and CB0205. Empty vector (EV) was used as a negative control to check for cross reactivity of the H6-antibody. The arrow marks the main protein (monomer) of interest.

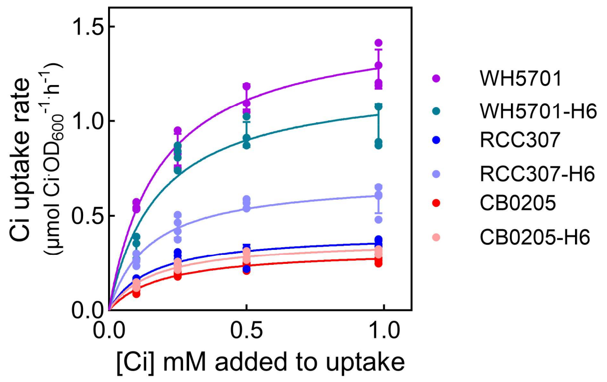


**Figure S4: $HCO_3^-$ uptake kinetics of tagged and untagged forms of SbtA2.** The affinity for $HCO_3^-$ was estimated using Ci uptake assays performed in response to increasing exogenous $HCO_3^-$ concentrations in *E. coli* DH5α in the presence of 20 mM NaCl. $HCO_3^-$ uptake rates at each $HCO_3^-$ concentration had empty vector rates specific for that condition subtracted. Similar trends were observed across experiments although the activity values varied, n=3 biological replicates, as a result representative data from single experiments are shown. Individual data values are shown along with their fitted Michaelis-Menten curve which is presented as a solid line. Curve fitting protocols are stated in the materials and methods section. Assay buffer composition: 20 mM BTP - $H_2SO_4$ pH 7.5 + 20 mM NaCl.

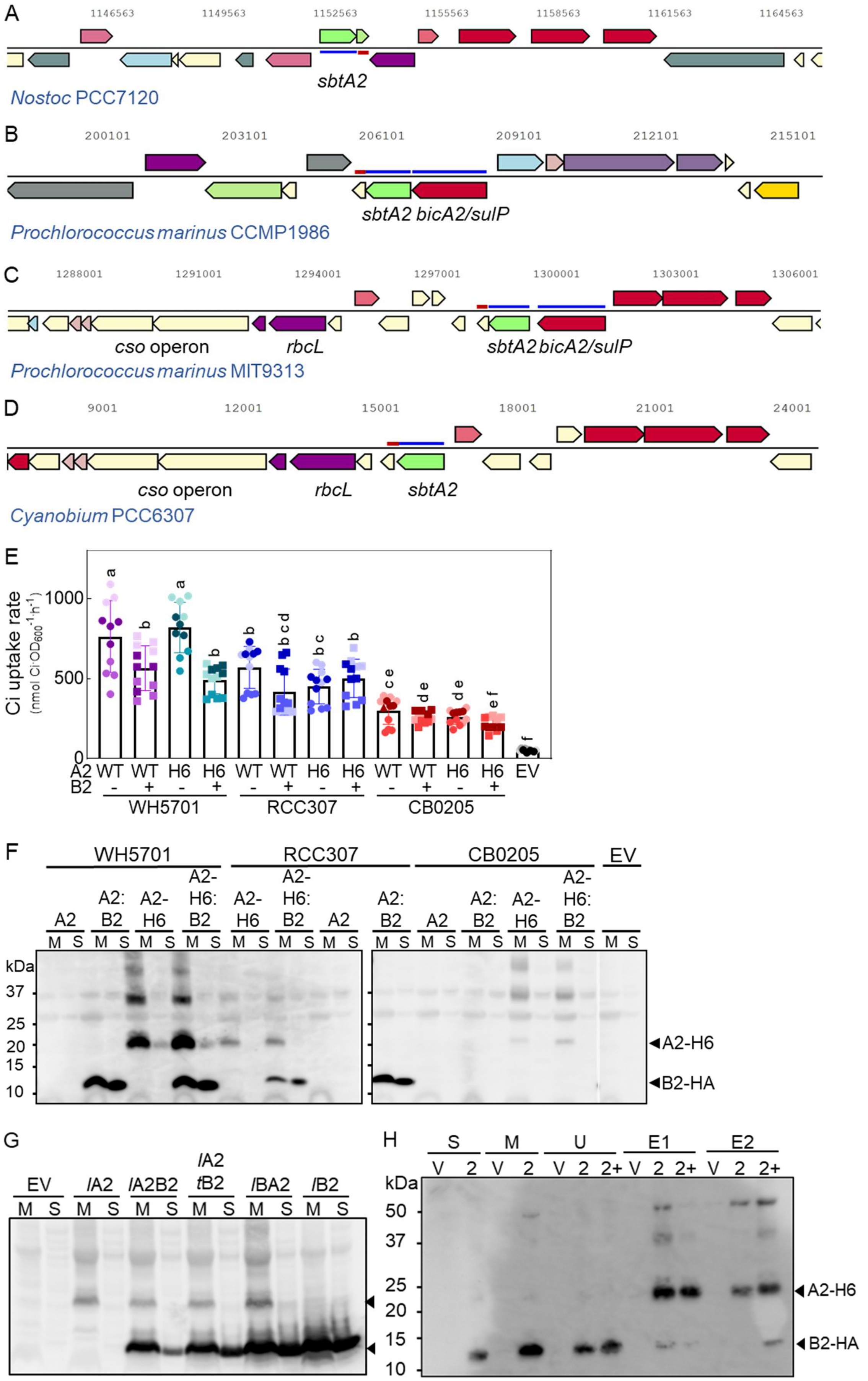
A
1146563
1149563
1152563
1155563
1158563
1161563
1164563
sbtA2
Nostoc PCC7120
B
200101
203101
206101
209101
212101
215101
sbtA2 bicA2/sulP
Prochlorococcus marinus CCMP1986
C
1288001
1291001
1294001
1297001
1300001
1303001
1306001
cso operon
rbcL
sbtA2 bicA2/sulP
Prochlorococcus marinus MIT9313
D
9001
12001
15001
18001
21001
24001
cso operon
rbcL
sbtA2
Cyanobium PCC6307
E
Ci uptake rate
(nmol Ci·OD600-1·h-1)
1000
500
0
A2
B2
WT
H6
EV
WH5701
RCC307
CB0205
F
WH5701
RCC307
CB0205
EV
A2
A2: B2
A2- H6
A2- H6: B2
M
S
kDa
37
25
20
15
10
A2-H6
B2-HA
G
EV
lA2
lA2B2
lA2 tB2
lBA2
lB2
M
S
H
S
M
U
E1
E2
V
2
2+
kDa
50
37
25
20
15
10
A2-H6
B2-HA

**Figure S5: SbtB2 gene arrangements and SbtB2 co-expression effects on SbtA2 $HCO_3^-$ uptake in *E. coli*.**

**(A-D)** Analysis of the genome context surrounding sbtA2 across different species of cyanobacteria.

*sbtA2* (shown in green with blue line) is typically observed with a small partner protein, likely *sbtB2* (shown with a red line) and sometimes in close proximity with other *CCM* genes. **(A)** *sbtA2* and *sbtB2* are localised together, as seen in *Nostoc* 7120. **(B)** *sbtA2* and *sbtB2* are located near bicA2 in some genomes, as seen in *P. marinus* CCMP1986. **(C)** *sbtA2* and *sbtB2* is located near *bicA2* and the *cso* operon, as seen in *P. marinus* CCMP9313. **(D)** *sbtA2* and *sbtB2* are in close proximity to the *cso* operon, as seen in *Cyanobium* PCC6307. Genome context surrounding *sbtA2* was gathered from Joint Genome Institute - Integrated Microbial Genomes and Microbiomes (Chen et al., 2022b)

**(E-F)** Co-expression of SbtB2 with SbtA2 and the effect on $HCO_3^-$ uptake function of SbtA2.

**(E)** $HCO_3^-$ uptake assays of *E. coli* expressing SbtA2±H6±B2-HA for WH5701, RCC307 and CB0205 forms. Data presented as mean ± SD, individual points represent means of an assay; n=3. Assay buffer composition: 20 mM BTP-HCl pH 7.5. Different letters indicate significantly different values when analysed by one-way ANOVA ($P<0.05$). **(F)** Western blot analysis of crude membrane-enriched and soluble fractions prepared from *E. coli* expressing of SbtA2±H6±SbtB2-HA from WH5701, RCC307, and CB0205 probed with anti-H6 and anti-HA antibodies. Untagged forms of SbtA2 and empty vector control used to ensure there is no cross reactivity of antibodies with the untagged proteins in the *E. coli*.

**(G)** *Varying the promoter on SbtA2 and SbtB2 altered the expression levels of these proteins* Immunoblot analysis of crude membrane enriched and soluble protein fractions prepared from DH5α *E. coli* carrying *lacI$^q$*-SbtA2-H6-WH5701 (*l*A2), *lacI$^q$*-SbtA2-H6:B2-HA-WH5701 (*l*AB2), *lacI$^q$*-SbtA2-H6:*tet*-B2-HA-WH5701 (*l*A2*t*B2), *lacI$^q$*-SbtB2-HA:A2-H6-WH5701(*l*BA2), *lacI$^q$*-SbtB2 (*l*B2) and a negative control (empty vector; EV). Blot probed with anti-HA and anti-H6 antibodies. Arrows indicate proteins of interest

**(H)** Pull-down binding assays to assess SbtA2-B2 complex formation.

Crude soluble (S) and membrane-enriched (M) fractions were made from E. coli co-expressing SbtA2-H6-SbtB2-HA. The fractions were then re-combined and incubated with $Ca^{2+}$ ± AMP before applying onto Profinity nickel beads. The H6-tagged moiety should bind to the column, including any proteins bound to it. Immunoblots of samples from the negative control (empty vector; V) and SbtA2-H6-SbtB2-HA (2) SbtA2-H6-SbtB2-HA + AMP (2+) were collected from the soluble (S), membrane-enriched (M), unbound (U), and two elution (E1, E2) fractions and were probed with anti-H6 and anti-HA antibodies.

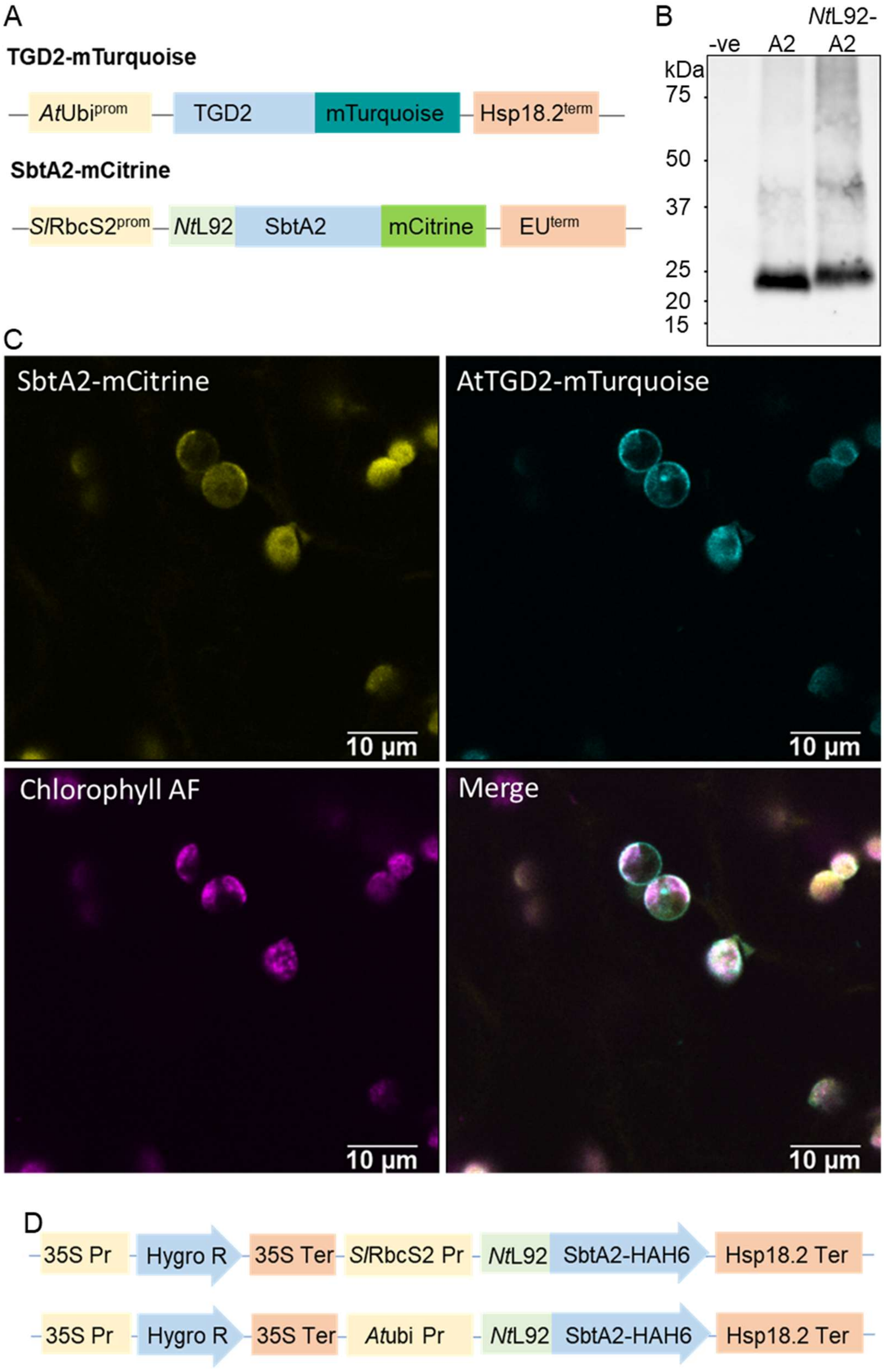
A
TGD2-mTurquoise
AtUbi prom
TGD2
mTurquoise
Hsp18.2 term
SbtA2-mCitrine
SlRbcS2 prom
NtL92
SbtA2
mCitrine
EU term
B
-ve
A2
NtL92-A2
kDa
75
50
37
25
20
15
C
SbtA2-mCitrine
AtTGD2-mTurquoise
Chlorophyll AF
Merge
10 µm
D
35S Pr
Hygro R
35S Ter
SlRbcS2 Pr
NtL92
SbtA2-HAH6
Hsp18.2 Ter
Atubi Pr

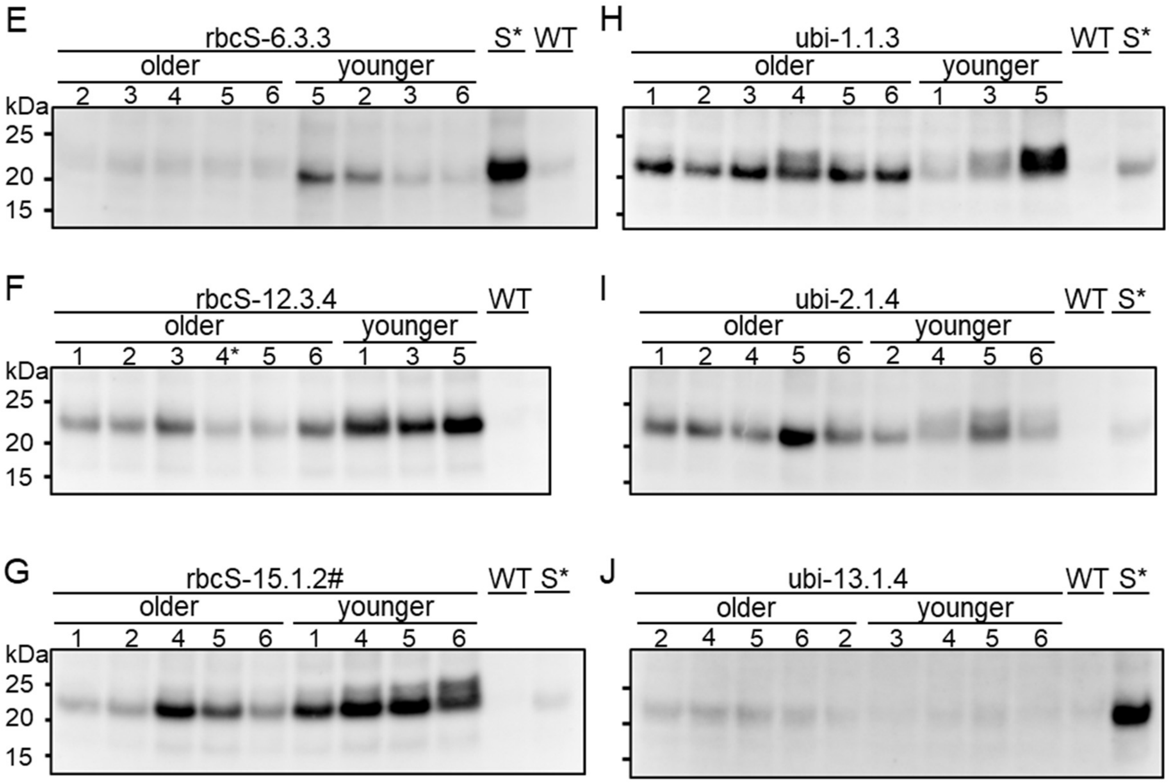


**Figure S6: Assessment of the localisation and expression of SbtA2 in *N. tabacum.***
**(A - C)** Assessment of the localisation of *Nt*L92-SbtA2 transiently expressed in *N. benthamiana.*
**(A)** Diagram of the constructs used for localisation of transiently express *Nt*L92-SbtA2-mCitrine and the IEM marker (DGD2-mTurquoise) in *N. benthamiana*. **(B)** Cleavage of the cTP from SbtA2 was determined by immunoblot analysis of membrane enriched samples prepared from *N. benthamiana* with transiently expressed SbtA2-HAH6 (C-terminally tagged) or *Nt*L92-PLGG1-SbtA2-HAH6. Blot was probed with anti-HA antibody. **(C)** Fluorescence microscopy of purified chloroplasts to identify the location of transiently expressed NtL92-SbtA2-mCitrine co-expressed with DGD2-mTurquiose (a chloroplastic IEM marker). The chlorophyll autofluorescence (AF) is also shown. The individual fluorophores were imaged and merged to determine if the two proteins co-located. **(D)** Two transformation constructs, differing in the promoter used to express SbtA2-HAH6-WH5701, were designed to transform into *N. tabacum* by *Agrobacterium*-mediated nuclear transformation.
**(E-J)** Assessment of SbtA2-HAH6-WH5701 protein expression in $Tob^{SbtA2}$ $T_2$ plants.
Immunoblot analysis of membrane-enriched fractions prepared from young and older plants of each $Tob^{SbtA2}$ line, and included S*, sample 4 from $Tob^{SbtA2\_RbcS\text{-}12.3.4}$, on each blot as a cross-sample control. Blots were probed with anti-HA antibody. Samples were taken from 5-10 cm in rosette size (younger) and 30 ± 5 cm (older) plants. Samples were loaded on an equal total protein basis. S* is sample 4 from $Tob^{SbtA2\_RbcS\text{-}12.3.4}$, included on each blot as cross-sample control. $Tob^{SbtA2\_rbcS\text{-}15.1.2\#}$ was not homozygous (G).

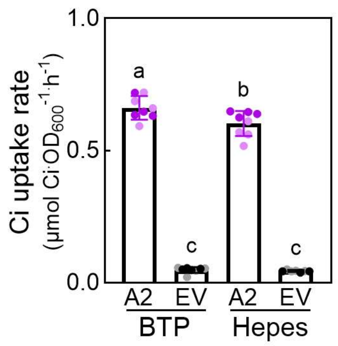


**Figure S7: Analysis of the basal $HCO_3^-$ uptake rate of SbtA2**

The $HCO_3^-$ uptake rates of DH5α carrying SbtA2-WH5701 (A2) or pCK1 (EV) using 2 different assay buffers, containing; 20 mM BTP-$H_2SO_4$ pH 7.5 or 20 mM Hepes-KOH pH 7.5. Data presented as mean ± SD; n=7-8 from 2 biological replicates. Different letters indicate significantly different values when analysed by one-way ANOVA ($P<0.05$).

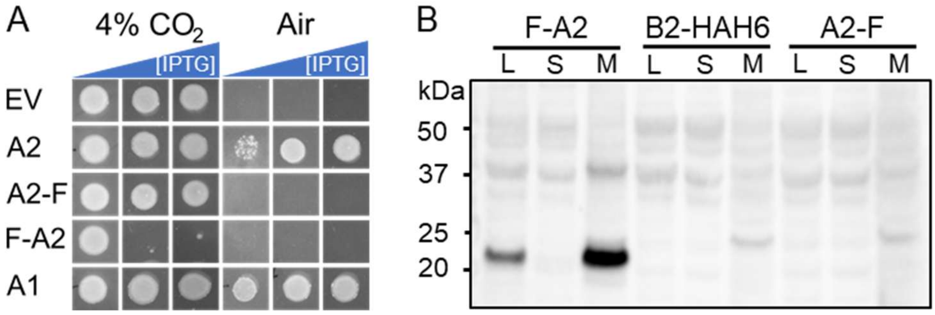


**Figure S8: Assessment of N- and C- terminally tagged SbtA2 $HCO_3^-$ transport function and protein expression.**

**(A)** Cultures of CAfree carrying SbtA1-7942 (A1), SbtA2 (A2), Flag-SbtA2 (F-A2), and SbtA2-Flag (A2-F) and empty vector were adjusted to the same density ($OD_{600}$) prior to spotting onto LB-agar and grown for 20 h at 37°C at ambient and 4% $CO_2$ under induction at 0, 50 and 200 µM IPTG. **(B)** Lysates (L), and crude membrane-enriched (M) and soluble (S) fractions were prepared from DH5α carrying Flag-SbtA2 (F-A2), SbtA2-Flag (A2-F) and SbtB2-HAH6 (B2-HAH6). The immunoblot was probed with anti-Flag antibody to detect the proteins. SbtB2-HAH6 was used as a negative control to check for antibody cross-reactivity. The constructs were expressed under the control of the *lacI$^q$-trc* promoter induced for 3 h with 1 mM IPTG.
